# How talin allosterically activates vinculin

**DOI:** 10.1101/2022.08.01.502287

**Authors:** Florian Franz, Rafael Tapia-Rojo, Sabina Winograd-Katz, Rajaa Boujemaa-Paterski, Wenhong Li, Tamar Unger, Shira Albeck, Camilo Aponte-Santamaria, Sergi Garcia-Manyes, Ohad Medalia, Benjamin Geiger, Frauke Gräter

## Abstract

The talin-vinculin axis is a key mechanosensing component of cellular focal adhesions. How talin and vinculin respond to forces and regulate one another remains unclear. By combining single molecule magnetic tweezer experiments, Molecular Dynamics simulations, actin bundling assays, and adhesion assembly experiments in live cells, we here discover a two-ways allosteric network within vinculin as a regulator of the talin-vinculin interaction. We directly observe a maturation process of vinculin upon talin binding which reinforces the binding to talin at a rate of 0.03 s^−1^. This allosteric transition can compete with force-induced dissociation of vinculin from talin only at 7-10 pN. Mimicking the allosteric activation by mutation yields a vinculin molecule that bundles actin and localizes to focal adhesions in a force-independent manner. Hence, the allosteric switch confines talin-vinculin interactions and focal adhesion build-up to intermediate force levels. The ‘allosteric vinculin mutant’ is a valuable molecular tool to further dissect the mechanical and biochemical signalling circuits at focal adhesions and elsewhere.

## Introduction

Tissue cells can sense, decode and respond to mechanical cues that depend on the physical and mechanical properties of their immediate surroundings, such as density, topography, and stiffness^1, 2^. Focal adhesions play a major role in the mechano-sensing capabilities of these cells as they develop when the tiny, dot-shaped and short-lived nascent adhesions inside the lamellipodia^3^. Myosin-II driven contractility grows these nascent adhesions into mature focal adhesions (FAs) and triggers a hierarchical recruitment of FA proteins which form large and complex cytoskeletal structures^4, 5^. The core regulatory action of these force-sensitive signalling hubs depends on an intricate interplay between talin, vinculin, and F-actin^3^.

Mechanical force plays a pivotal role in all stages of focal adhesions development and turnover. The forces within the cell – generated to a large extent by the actomyosin machinery – can fluctuate strongly in location and time. Interestingly, the locations that exhibit the highest cellular forces have been identified to often coincide with the assembly sites of FAs^6, 7^. Tension sensors indicated that talin and vinculin experience forces as high as 10, pN when engaged in FAs^8–10^. Within FAs, vinculin acts as a mechanical reinforcement in the link between the actin cytoskeleton and talin, which in turn is directly bound to the integrin receptors connecting to the celĺs exterior^4, 10, 11^.

Vinculin, a 5-domains protein composed of 1066-residue-long, is present in all force-bearing cellular junctions such as FAs, adherens junctions, and immunological synapses^12–14^. As an interaction hub and versatile signalling molecule, vinculin harbours binding sites for a variety of adhesome proteins, present in these distinct junctional systems^15^. In its auto-inhibited conformation, vinculin’s tail domain (Vt) packs onto its head domain (Vh), and most of its binding sites are cryptic. For vinculin to unfold its full signalling potential, activation is required^16^. The exact process by which vinculin is activated is not yet fully understood.

The head domain Vh comprises four helix bundles, D1-D4, with the D1 domain harbouring the interaction site for talin as well as for α-actinin, and α-catenin. Their so-called vinculin binding sites (VBS) are single helical domains, which are packed up in helix bundles and masked if mechanical force is absent^17, 18^. Talin contains eleven VBS; α-actinin and α-catenin only contain one VBS^19–21^. The structural mechanism by which VBS binds to vinculin is conserved across all of vinculin binding partners^20, 22, 23^. X-ray crystallography structure determination of D1-Vt with and without a bound VBS revealed a mechanism in which VBS binding reorganizes the helix bundle of D1 such that vinculin is partially activated^23^.

The three known actin binding sites on the vinculin tail are either partially or fully occluded in the auto-inhibited conformation and, hence, actin is not able to bind to on its own^24–26^. Binding of actin to full-length vinculin requires an artificially weakened head-tail interface^27^ or VBS binding^28^. Bois et al.^20^ suggested that the α-actinin VBS alone can cause vinculin activation, while other studies propose a collaborative effort of a VBS and actin^29, 30^. Further players in the multimodal activation scenario of vinculin include mechanical force^31^ and PIP2^32, 33^.

While mechanical force across the talin backbone has been unequivocally shown in single molecule studies, to be decisive for VBS exposure and vinculin binding^34–36^, a direct role of force for vinculin activation is less apparent. A collision-induced semi-open vinculin state with an increased cross-section area was recently captured using ion-mobility mass spectrometry^37^, possibly mimicking a force-induced activated state of vinculin. Interestingly, decreased cellular traction forces increase the unbinding of zyxin from FAs but not of vinculin^38^. Recent findings by Boujemaa-Paterski et al.^28^ revealed an enhanced actin-bundling by vinculin in a force-independent manner, that is, which takes place when full-length vinculin is exposed to a talin VBS1 domain. These findings support the hypothesis that VBS-binding activates vinculin – being it fully or partially – and establish an order of events in which VBS binding proceeds actin binding and thus force transmission onto vinculin.

To test this hypothesis, we used a multidisciplinary integrative approach comprising Molecular Dynamics simulations, single molecule force spectroscopy experiments, functional in vitro assays, and live cell imaging approaches. With these tools, we reveal a two-way allosteric mechanism in full atomistic detail of how vinculin regulates talin binding and vice versa. On this basis, we rationally designed vinculin mutants that aim to mimic the underlying destabilisation in the head-tail auto-inhibitory interface by VBS. For our designed vinculin mutants that simulate the VBS effect on vinculin, we tested actin bundling in vitro and focal adhesion formation in live cultured cells, both in a largely force-independent fashion. Our study disentangles the key steps of vinculin activation and identifies an extensive rewiring of a salt-bridge network as the key mechanism in weakening vinculin’s auto-inhibitory head-tail interface. It also puts forward our vinculin mutants as close mimics of vinculins bound to stretched talin and alpha-actinin, and proposes them as novel molecular tools to study and interfere with cell-matrix and cell-cell adhesion.

## Results

### Vinculin’s head-to-tail interaction weakens vinculin-talin binding

Vinculin can bind talin in its active and autoinhibited state. But if vinculin in its active open state binds to talin any different than its inactive closed state is unknown. We investigated the head-to-tail interactions in vinculin impact talin binding by force-probe MD simulations. We subjected VBS1 to a set of different constant forces in the range of 200-260pN (**Fig. 1A-C**), while being bound to either full-length vinculin (FLV) or only to vinculin’s D1 domain (Vd1). We started from a VBS1-vinculin complex (see Methods) and monitored the progression of VBS helix force-induced unravelling by the increase of end-to-end distance (**Fig. 1B, Fig. S1**). We then deduced Mean First Passage Times (MFPT) for the transition to the first partially unfolded intermediate of VBS (**Fig. 1C**). Across the force regime covered in the simulations, VBS1 shows, on average, shorter MFPTs **(Fig. S1D,E)** by approximately one order of magnitude when bound to FLV compared to Vd1, while the force-dependence of these times is comparable (linear Bell-Evans fits in **Fig. 1C**). This suggests the presence of the head-to-tail interaction to destabilize the complex, leading to a faster loss of helicity and vinculin-talin interactions under force. The atomistic detail of the simulations allowed the identification of a range of residues within the vinculin D1 domain that VBS1 and the tail compete for when bound to vinculin **(Fig. S2)**, explaining this allosteric mechanism.

**Figure 1:**
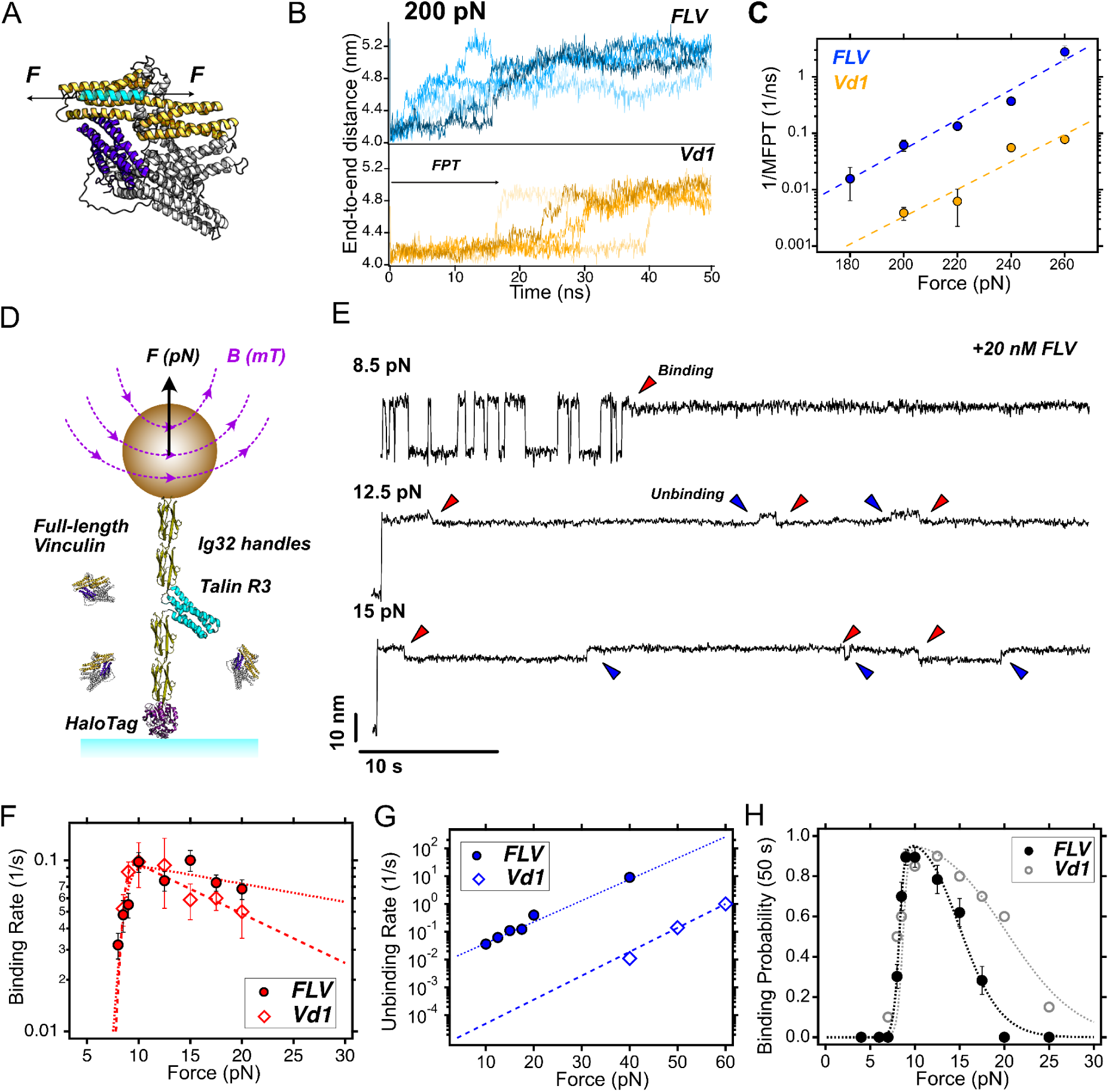
Vinculin’s head-tail interface regulates vinculin-VBS binding. **(A)** Structural scheme of full-length vinculin (FLV). D1 domain (Vd1) shown in orange, and tail domain in purple binding to a VBS under force (cyan helix). **(B)** End-to-end length at 200 pN of VBS1 bound to full-length vinculin (upper panel) and vinculin D1 domain (lower panel) as observed in MD simulations. The increase in extension from 4.0 nm to 5.2 nm corresponds to the first uncoiling transition of VBS1, which we use as a proxy for vinculin unbinding (first passage time, FPT, as indicated). VBS1 bound to FLV unbinds much faster than when bound to Vd1. Data for other forces shown in Fig. S1. **(C)** Expected rates (inverse MFPT, computed as shown in Figs S1D,E) to first transition (vinculin unbinding) as a function of force for FLV (blue) and Vd1 domain (orange). Data fitted to a Bell-Evans’s model (FLV: k_0_=2.6×10^−7^ s^-1^, x^†^=0.25 nm; Vd1: k_0_=4.5×10^−8^ s^-1^, x^†^=0.23 nm). **(D)** Single-molecule magnetic tweezers assay to measure full-length vinculin binding to talin under force. **(E)** Magnetic tweezers trajectories of R3^IVVI^ in presence of 20 nM full-length vinculin. At 8.5 pN (upper panel), talin folds and unfolds in equilibrium, exposing its vinculin binding sites. Single full-length vinculin binding events can be detected as a ∼3 nm contraction of the talin polypeptide (red arrow) and the stopping of talin’s folding dynamics. At higher forces (12.5 pN, middle panel, and 15 pN, lower panel), the vinculin binding sites are fully exposed and FLV quickly binds. However, unlike what previously reported for the Vd1 domain, the complex is much less stable and FLV unbinds after a few seconds, showing an upward ∼3 nm step (blue arrow). **(F)** Binding rates of FLV and Vd1 to R3^IVVI^. The binding kinetics of FLV and Vd1 are comparable, showing first a steep increase at ∼8 pN associated with R3 unfolding rates, and then a decrease with force, due to the energetic penalty of the coil-to-helix transition triggered by vinculin binding. Data is fitted to a double-Bell’s model, with a positive and negative force dependency. **(G)** Unbinding rates of FLV and Vd1. The unbinding rates of FLV are much faster than those of Vd1, indicating a less stable complex. (FLV: k_0_=6.6×10^−3^ s^-1^, x^†^=0.72 nm; Vd1: k_0_=6.8×10^−6^ s^-1^, x^†^=0.81 nm). **(H**) Binding probability of FLV and Vd1 measured over a 50 s time window.

We next tested this prediction experimentally. To measure the interaction between FLV and talin under force, we purified the proteins and applied single-molecule magnetic tweezers to directly detect individual vinculin-binding events to single talin molecules subjected to forces in the picoNewton (pN) range. Previously, we used magnetic tweezers to characterize the force-dependent binding of Vd1 to the talin R3 domain^35^. These experiments showed that force regulates Vd1 binding in a biphasic way; force first favors Vd1 binding by unfolding talin and exposing the cryptic VBSs, but above 20 pN this interaction is hampered due to the coil-to-helix on talin’s VBSs transition triggered by vinculin binding, which further provides an unambiguous fingerprint to detect individual vinculin binding events. Based on that assay, we here investigated the binding of FLV to single talin R3 domains subjected to pN-level forces. In our experiments, the R3^IVVI^ domain—the R3 domain harboring the IVVI mutation, which increases its mechanical stability without impacting vinculin binding^35, 39, 40^—is flanked by two mechanically stiff Ig32 domains, serving as molecular handles (**Fig. 1D**). At the C-terminus, a HaloTag allows specific and covalent anchoring to a glass coverslide, while a biotinylated N-terminus AviTag binds to streptavidin-coated M270 superparamagnetic beads, establishing a highly specific and stable single-molecule tether. By controlling a magnetic field using a magnetic tape head^41, 42^, we can subject single R3 domains to low mechanical forces (0-40 pN) and monitor its conformational dynamics over physiologically-relevant time scales. (**Fig. 1D**).

In the presence of 20 nM FLV, our experiments indicate that FLV binds talin R3 in the same way as Vd1, contracting the unfolded talin polypeptide by ∼3 nm and locking talin in the unfolded state (**Fig. 1E**, red arrow), which suggests that both FLV and Vd1 bind talin through an analogous mechanism. However, while at low forces—*e.g.* 8.5 pN, where talin folds and unfolds in equilibrium—no apparent differences between FLV or Vd1 binding are noticed (**Fig. 1E**, upper panel), when we explored higher forces such as 12.5 pN or 15 pN (**Fig. E**, middle and lower panel), we observed that FLV unbinds over just a few seconds, showing an upwards ∼3 nm step that indicates the uncoiling of the VBSs and the recovery of talin’s unfolded extension (**Fig. E**, blue arrows). This is different from the case of Vd1, which forms a highly stable complex, requiring forces in excess of 40 pN to trigger vinculin unbinding over seconds-long timescales^35^.

To characterize the nanomechanics of the R3-FLV complex and compare them to those previously determined for Vd1, we measured the force-dependent binding (**Fig. 1F**) and unbinding (**Fig. 1G**) kinetics of FLV to talin R3^IVVI^. Interestingly, the binding rates are equivalent to those previously measured for Vd1, which suggests that the binding reaction (on-rate) is dominated by the vinculin D1 domain alone, with no or little influence of the rest of vinculin head domains (D2-D4), or the tail domain. The biphasic force dependence of the binding rates can be modelled by a double-exponential Bell model, where the first increase of the binding rates with force is governed by talin unfolding rates, while the decrease at higher forces arises due to the energetic penalty of reforming the VBSs helices under force. By contrast, the unbinding rates of FLV are very different from those measured for Vd1. Similar to the behaviour observed in the MD simulations (**Fig. 1C**), the slope of the force dependence is very similar in both cases (FLV: x^†^=0.72 nm and Vd1: x^†^=0.81 nm), which indicates a comparable distance to transition state in both cases, likely dominated by the uncoiling of the VBSs. However, the intrinsic off-rate differs by three orders of magnitude (FLV: k_0_=6.6×10^−3^ s^-1^, and Vd1:k_0_=6.8×10^−6^ s^-1^), indicating that full-length vinculin forms a much less stable complex with talin R3 than the vinculin D1 domain. When characterizing the equilibrium binding properties as the binding probability measured over a 50 seconds time window (**Fig. 1H**), FLV shows a much narrower force-range for binding, being optimal between forces of 8 and 15 pN. Overall, our MD simulations and single-molecule experiments together indicate that full-length vinculin forms a less stable complex with talin than the isolated D1 domain, suggesting that, while auto-inhibited vinculin is still capable of binding talin under force, the head-tail vinculin interaction allosterically renders a weaker complex that unbinds after only a few seconds. A stable complex to be formed likely requires vinculin activation by the release of the head-tail interaction, which we investigated next.

### VBS binding facilitates vinculin activation in MD simulations

As suggested by crystallographic data, VBS binding causes major structural changes in the vinculin D1 domain^23^. To reveal the implications of these changes on the strength of the D1-tail interface in full-length vinculin and to rationally design mutants to interfere with these changes, we subjected the equilibrated VBS1-vinculin complex and the apo-state protein to force-probe MD simulations. We now employed an external force between the head and tail domain to enhance the sampling and provoke a dissociation along the most likely pathway of *in vivo* vinculin activation, on the time scale of the MD simulations. **Fig. 2** depicts the simulation system of the VBS1-vinculin complex, with snapshots extracted from the trajectories (snapshots for the apo-state of vinculin are shown in **Fig. S3** for comparison).

**Figure 2:**
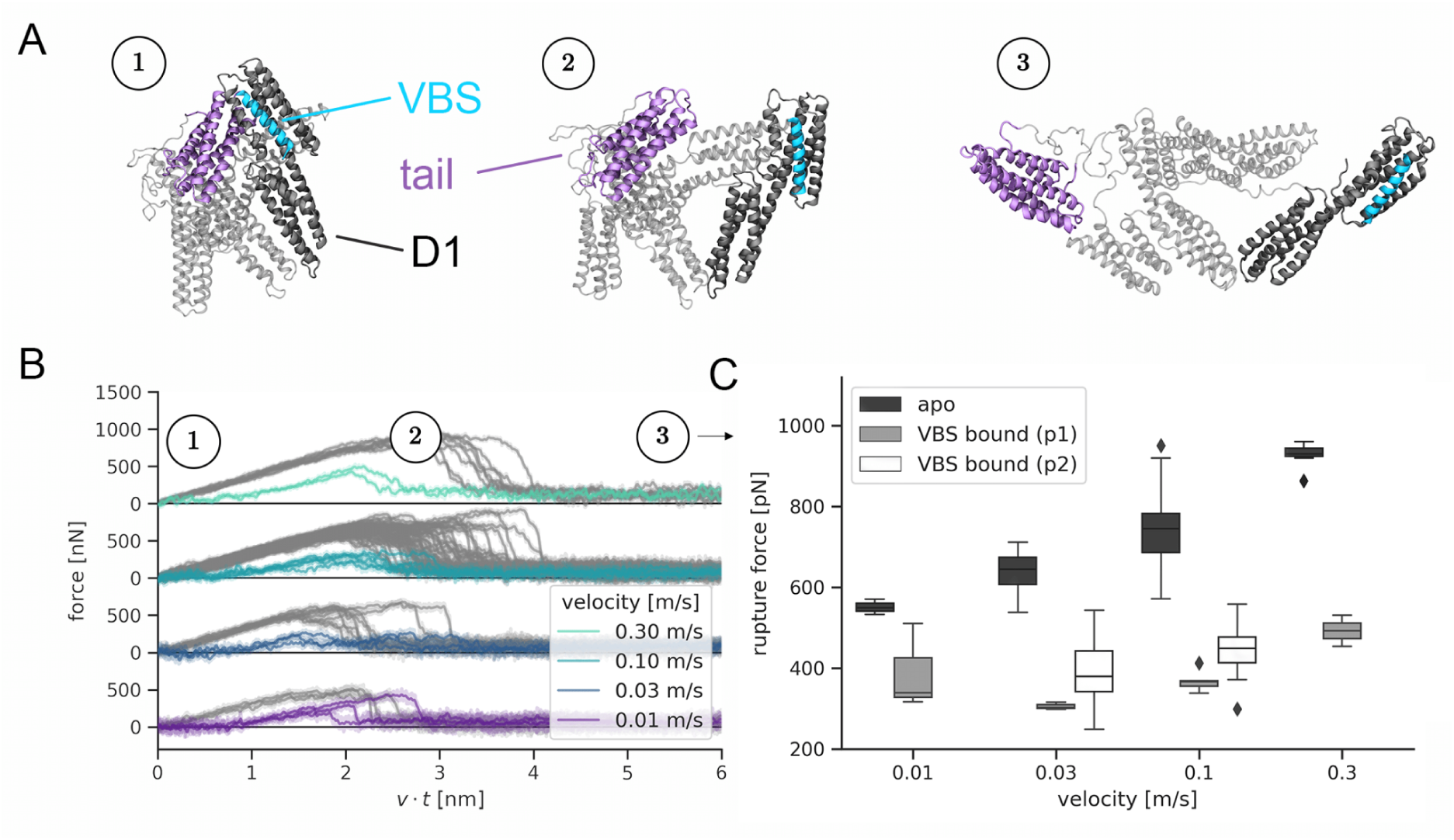
Talin binding facilitates vinculin head-tail opening. **A)** Snapshots extracted from an exemplary trajectory to represent the opening transition during the force-probe MD simulations (see **Figure 1B** and **Methods**). Force was applied directly to the VBS peptide (cyan) or to the talin binding site located on the vinculin D1 domain (dark grey). All simulations included full-length vinculin (Vh in grey, Vt in purple) **B)** Force-extension traces for force-application to VBS1 and pulling speeds between 0.01 and 0.3 m/s. As a reference, the force-extension curves for opening of the apo-state protein are shown in grey. **C)** Quantification of the recorded rupture forces for the apo-state. For the VBS-complex rupture forces were decreased drastically and two points of force application were investigated: (p1) on vinculin at the location of the talin binding site (light grey), which ensued slightly higher rupture forces, and (p2) directly on the VBS (black), which lead to the smallest observed rupture forces.

As expected, the tail domain (purple in **Fig. 2A**) detaches from the head domain (dark grey). All helical bundles remain intact, and the VBS1 peptide (cyan) stays stably bound to D1. The simulation protocol thus successfully samples the transition towards an activated vinculin conformation. The force-extension curves (**Fig. 2B**) reveal a pronounced drop in the force required for detaching the tail from the remainder of the protein bound to VBS1. In the apo-state of vinculin, where VBS1 is absent, force-extension curves reach pronouncedly higher rupture forces for opening the head-tail interface. **Fig. 2C** directly compares the rupture forces as a measure for the strength of the auto-inhibitory head-tail interface across four different pulling velocities. VBS1 binding consistently lowers, that is, roughly halves, the rupture force for vinculin activation, e.g. from ∼550pN to ∼350pN at the lowest pulling velocity of 0.01m/s. Additional simulations excluded that the observed drop in rupture forces is caused by the slight difference in pulling direction (pulling from VBS helix versus talin binding site, for details see **Fig. 2C** and Methods). To ensure comparability, we limited subsequent analysis and simulations to the case in which force is consistently applied to the talin binding site.

To further validate that the pronounced drop in resistance against activation of vinculin is solely due to the presence and entailed conformational change of VBS and does not arise from other differences in the crystallographic starting structures, we initiated an additional set of simulations from an apo state obtained by deletion of VBS1 from the complex. Remarkably, we observed both the recovery of the D1 conformation close to the experimentally determined apo state as well as of the high activation forces characteristic for this state (**Fig. S4**).

### Talin Binding Induces a Partial Dissociation of Vinculin’s Head-Tail Interaction

The computational evidence suggests that talin binding facilitates vinculin activation by allosterically weakening the head-tail interaction. Therefore, we wondered whether such conformational transition in vinculin upon binding to talin under force could be captured with our single-molecule approach. While in our magnetic tweezers experiments we monitor the conformational dynamics of talin and not of vinculin, our experiments strongly suggest that the head-tail interaction weakens the interaction of vinculin with the VBSs under force. Thus, it is enticing to hypothesize that, if vinculin activation occurs upon binding to talin as an opening (or at least a partial opening) of the head-tail interface, such conformational change would be indirectly measurable at the single-molecule level as an increase in the stability of the talin-vinculin complex under force.

When measuring R3 dynamics at low forces (∼8.5 pN) in the presence of FLV over several hours, we observed that vinculin unbinds over a timescale of an ∼hour (**Fig. 3A**). These data indicate that, under physiological forces, FLV forms a stable complex with R3, compatible with the lifetime of focal adhesions^43, 44^. However, these slow unbinding kinetics are not consistent with those shown in **Fig. 1G**, which, extrapolated to forces around 8.5 pN, imply much faster unbinding rates (∼0.01 s^-1^). Thus, this seemingly contradictory evidence, suggests the existence of two different bound states with a different stability. To test this hypothesis, we designed a force protocol to examine the stability of the R3-FLV complex (**Fig. 3B**). First, we allowed FLV bind talin at 8.5 pN and kept the complex formed at this low force for a variable time to then interrupt it with a high-force 40 pN pulse, triggering the mechanical dissociation of vinculin. By applying the high force pulse at different moments after the binding event is observed, we characterized the stability of the complex as a function of the complex lifetime. When we applied the 40 pN pulse soon after vinculin binds, we observed a very rapid dissociation, in the sub-second timescale (**Fig. 3B**, left panel, inset); however, if the high-force pulse is applied several seconds after the R3-FLV complex is formed, the unbinding kinetics are much slower, taking a few seconds to dissociate (**Fig. 3B**, right panel). The existence of two different bound modes was confirmed when measuring the distribution of unbinding times at 40 pN, which shows a clear double-exponential behavior, with a first weak mode that dissociates over ∼0.4 seconds, and a second stronger one that unbinds over ∼7 s (**Fig. 3C**). To determine whether the existence of these two bound modes was related to the lifetime of the complex—perhaps involving a maturation process to render a tighter interaction—we calculated the unbinding time as a function of the complex lifetime (**Fig. 3D**). These data show a sigmoidal dependence, indicating that over very short timescales the interaction is characterized by a low stability (compatible with the unbinding rates measured in **Fig. 1G**), while after a few seconds, the complex evolves towards a more stable interaction, compatible with the slow unbinding rates measured at 8.5 pN (**Fig. 3A**). From this sigmoidal dependence, we can estimate the timescale for this maturation transition to be ∼37 s.

**Figure 3:**
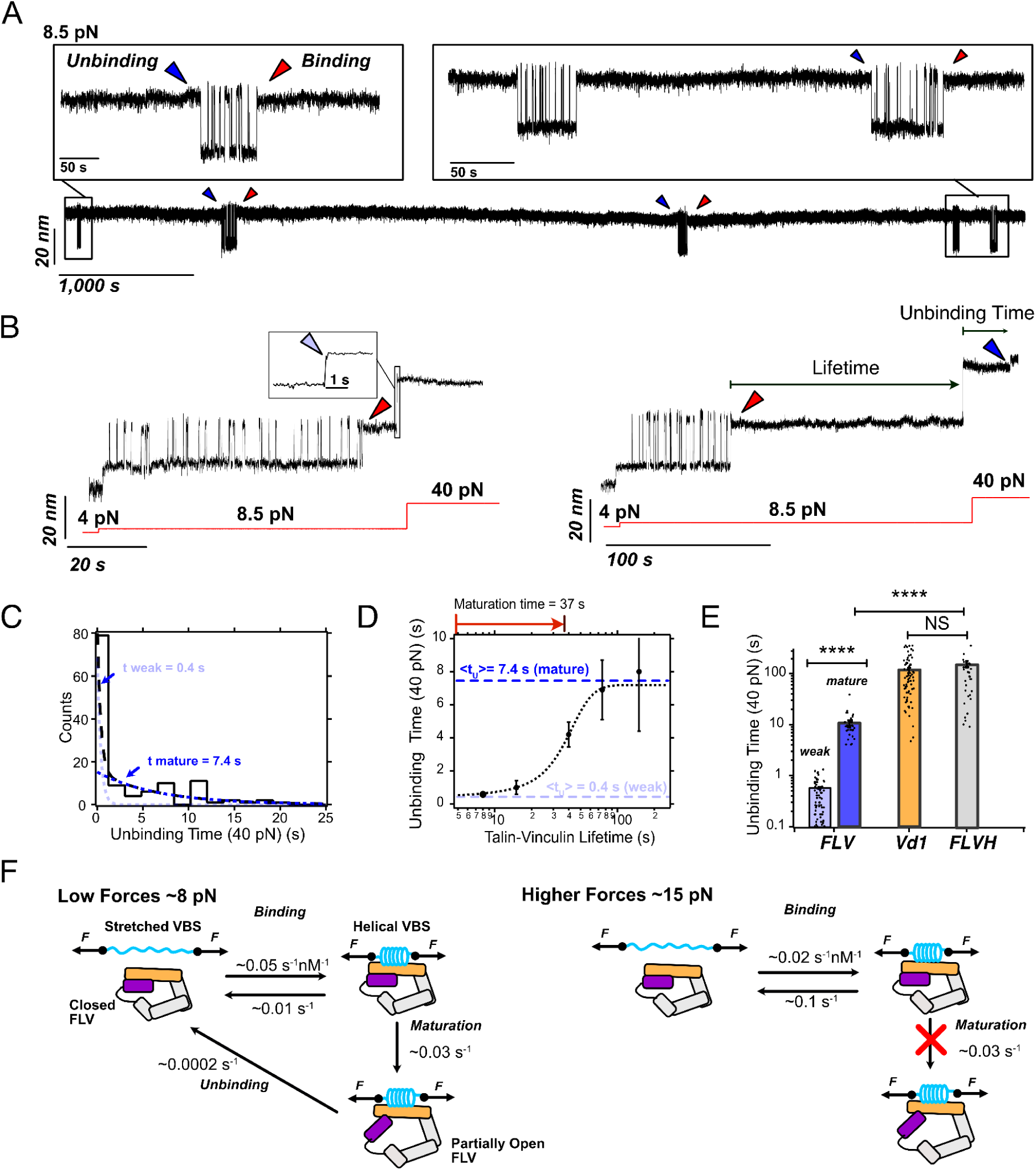
Talin binding weakens the vinculin head-to-tail interaction. **(A)** Magnetic tweezers recording showing the binding dynamics of full-length vinculin to talin R3^IVVI^ at low forces over hours-long timescales. At 8.5 pN, FLV can remain bound to talin over ∼1 h, indicating much slower unbinding kinetics than those corresponding to higher forces. **(B**) Magnetic tweezers recordings where the talin-vinculin complex is interrupted by a 40 pN high force pulse after bound for just a few seconds (left) or several seconds (right). Over short lifetimes, FLV unbinds on a sub-second timescale, while if the complex is left to mature at low forces, the interaction is reinforced, and it unbinds at 40 pN after a few seconds, indicating a much more stable complex. **(C)** Distribution of unbinding times (t_U_) at 40 pN. The unbinding kinetics show a double-exponential distribution, with a fast timescale of ∼0.4 s (weak) and a slower timescale of ∼7.4 s (mature), indicating two different binding modes. **(D)** Unbinding time of FLV at 40 pN plotted as a function of the complex lifetime. The talin-vinculin interaction matures towards a more stable complex over a timescale of ∼37 s. **(E)** Unbinding kinetics of FLV (weak state, light blue; mature state, dark blue), the vinculin D1 domain (orange), and full-length vinculin head (grey). **(F)** Kinetic diagram suggesting the binding mechanism of FLV to a VBS low force (left) and higher forces (right). At forces in the ∼8 pN range, the interaction is reinforced over ∼37 seconds, likely due a partial opening of vinculin that stabilizes the talin-vinculin complex. At higher forces around ∼15 pN, the maturation step is kinetically unfavourable due to the faster competing unbinding kinetics, and FLV only binds VBS on its weak mode.

Given that Vd1 forms a much more stable interaction with R3, also involving just a single bound state, we hypothesized that the observed maturation involves some conformational transition in FLV, likely related to vinculin activation. To this aim, we measured the binding of full-length vinculin head (FLVH, D1-D4 domains) to R3, which strikingly showed the same binding and unbinding properties as Vd1 (**Fig. 3E** and **Fig. S5**). This indicates that, from the vinculin head, only the D1 domain plays a role in interacting with talin. However, the dissociation kinetics of the mature state of FLV are still faster than those of Vd1 and FLVH, indicating the weakening effect of the vinculin tail domain still persists even on the mature bound state. This could suggest that talin alone is not sufficient to fully release the head-tail interface, and that the allosteric effect of this interaction renders a “pre-active” vinculin, perhaps requiring actin binding for full activation.

Thus, based on our single-molecule data, we propose a kinetic model for the vinculin-binding mechanism (**Fig. 3F**). At low forces (up to ∼9-10 pN), FLV binds the stretched VBSs with very high affinity (estimated Kd∼0.2 nM). Upon binding, vinculin undergoes a conformational transition (likely a partial opening of the head-tail interface) over a timescale of ∼37 s, that renders a much more stable complex (Kd∼0.004 nM). However, if talin is under higher forces above ∼12.5 pN (still within the physiological range), although vinculin still binds quickly with high affinity (Kd∼5 nM), the unbinding rates are too fast to allow vinculin maturation, rendering a weak complex that dissociates shortly after it forms.

### Rewiring of salt bridge network causes vinculin activation upon talin binding

Talin binding to vinculin strongly undermines vinculin’s resistance against opening of the auto-inhibitory head-tail interface, as shown by our MD simulations and magnetic tweezers experiments. While the comparison of the apo and VBS1-bound states already highlighted the role of electrostatic interactions^23^, the MD simulations now provide the full dynamic picture at atomistic detail and allow to pinpoint the key residue interactions allosterically affected by VBS binding. We calculated the major correlated motions present at the interface of Vh (residues 1-200) and Vt (880-1066) in the tensed state (**Figure 4A** and Methods). The apo state shows several clusters of strongly correlated interface residues with significantly diminished correlation upon VBS1 binding (**Figure 4B**). The residue pairs with the highest loss of correlation within each cluster (**Table S1**) are visualised as spheres in **Figure 4C** which also displays the networks of increased and diminished interactions as identified by force distribution analysis (see methods section). The overlap in results of the two analysis methods (one based on non-equilibrium simulations, the other on equilibrium simulations) allowed us to locate three areas of strong coupling: residues K944 and R945, as well as residues D1013 and E1015 give away most of their coupling to the regions between the tail residues 19-33, and 88-116. In addition, tail residue E986 loses interactions with a region around residue 185. On this basis, we designed two novel mutants, K944A-R945A-D1013A-E1015A (4M) and an additional mutation E986A (5M), in which the charged residues in the head domain are mutated to alanines in order to abolish the stabilising salt bridge network, thereby mimicking the VBS-bound matured state. We tested if the allosteric mechanism and its mimic by 4M and 5M are robust across different VBS, and to this end performed additional MD simulations with VBS3 and alpha-actinin (**Fig. S6**). VBS3 binding again caused a loss of correlations involving all those five residues, resembling 5M, while alpha-actinin binding did not affect E986 interactions, resembling 4M (**Fig. S6A**). In accordance, VBS3 binding measurably weakened vinculin head-tail interactions, while alpha-actinin did not show significant changes within the simulated time scale (**Fig. S6B**). We predicted these mutants, due to the weakened head-tail interactions, to show stronger binding to talin, that is, binding times more comparable to the vinculin head domain than to full-length vinculin.

**Figure 4:**
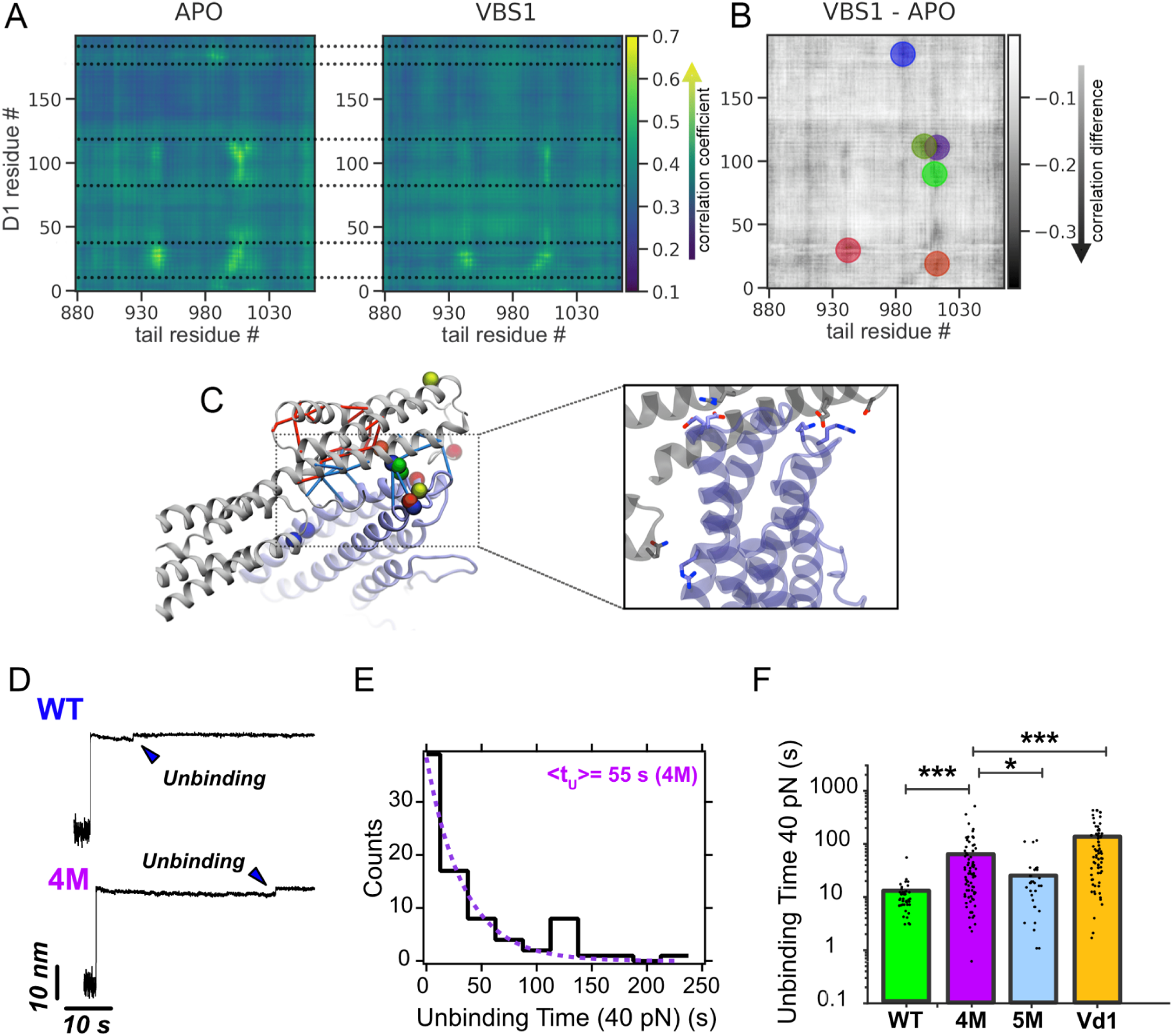
Analysis of correlated motions and force distribution identified an overlapping set of residues that collectively weakens the D1-tail interface. **A)** The results of the analysis of correlated motions^45^ in the tensed protein. (Extracted from force-probe MD trajectories at 0.1m/s using the 5 ns leading up to a force of 300 pN across the protein in each trajectory. For each map, the contributions from 20 independent trajectories were averaged.) Here, residues of the vinculin tail are represented along the x-axis, and residues of the D1 domain along the y-axis. The areas between the dotted lines illustrate the three areas in the D1 domain that are strongly affected by VBS-binding. **B)** Shows the changes in correlation coefficient per residue between APO state and complex, such that dark areas represent residues that experience a strong loss of correlation if in complex with VBS1. The colored circles represent the results of a weighted cluster analysis, intended to find clusters of correlation-losing residues **C)** Representation of the results from the cluster-analysis (colored spheres) and FDA-network analysis (red and blue connections). The zoom-in on the right shows the interactions between head and tail residues (image based on the x-ray structure) that were identified by both methods to contribute to the lesser stability of the head-tail interactions. **D)** Magnetic tweezers trajectories showing vinculin unbinding from talin R3^IVVI^ at 40 pN for the WT (upper panel) and 4M mutant (lower panel). The 4M mutant has much slower unbinding kinetics, indicating that it forms a more stable complex. **E)** Distribution of unbinding times for the 4M vinculin mutant. The unbinding times are distributed as a single-exponential, indicating that, unlike WT vinculin (compare Fig. 3C), the 4M mutant does not undergo any maturation process, at least within the experimentally accessible timescales. **F)** Unbinding times for WT vinculin (mature state, dark blue), 4M (purple) and 5 M (blue) mutant, and D1 domain (orange). The 4M and 5M mutants show much increased unbinding times, approaching those of the D1 domain, suggesting that the mutations weaken the D1-tail interface, which results in a more stable talin-vinculin complex.

To test this predicted effect of the designed mutations (4M and 5M vinculin mutants) on the association between talin and vinculin, we used our magnetic tweezers assay to measure the stability of the mutant vinculin-talin interaction. When forcing the complex dissociation at 40 pN (**Fig. 4D**), we readily observed that the unbinding kinetics of the 4M and 5M mutants from talin were slower than those measured for WT vinculin, even compared to the mature state. Additionally, the unbinding times for both mutants were distributed as a single-exponential, indicating a single binding mode (**Fig. 4E**). Similarly, the 5M mutant shows comparable properties to its 4M counterpart (**Fig. 4F**). While we cannot rule out that the mutant vinculins undergo an opening transition similar to that suggested for the WT, it must occur on a timescale faster than our experimental resolution (we typically need ∼5 seconds to ensure that vinculin binds), suggesting that the mutations severely weaken the D1-tail interface, facilitating the allosteric relationship between talin binding and vinculin activation. These data together indicate that the mutant vinculins form a stronger complex with talin, approaching to the stabilities measured for the isolated D1 domain (**Fig. 4F**), likely because the mutations weaken the head-tail interface leading to a pre-opened conformation.

### Efficient bundling of actin filaments by the vinculin 5M and 4M mutants

We next examined the functional consequences of the designed vinculin mutants 4M and 5M in vitro. Vinculin has been shown to form stable actin bundles in the presence of activated talin^28, 46^. Here, we used total internal reflection fluorescence (TIRF) microscopy to investigate the ability of the mutants to bundle actin filaments even in the absence of activated talin (**Fig. 5**). We monitored the polymerisation of fluorescently labelled actin monomers in the presence of only the full-length vinculin variants or in the presence of variable amounts of the talin constitutively active vinculin-binding site 1 (VBS1, residues 482–636)^22, 27, 28^ (**Fig. 5A** and **movies S1 and S5**). In the absence of talin-VBS1, the vinculin 5M variant induced stable actin bundles with a ratio of 40.3 ± 10.6 of the total population of filaments, 1.4 times greater bundles yielded by vinculin 4M variant (**Fig. 5B**). Surprisingly, the bundles formed in the presence of the vinculin 4M variant were less stable than those formed by the 5M variant (**movies S2**). Furthermore, in control assays we confirmed that the actin bundling in the absence of talin-VBS1 was due to the mutations in vinculin since the polymerisation kinetics of pure actin or supplemented with wild-type vinculin showed rarely and only momentarily actin filaments crossings (**Fig. 5C** and **movies S1 and S2**). This suggest that the mutations in the 4M protein, K944A, R945A, D1013A, E1015A, perturbed the head-to-tail interactions of vinculin and therefore induced actin bundling, while the additional mutation E986A significantly shifted vinculin dynamics towards a state prone to bind and stably bundle actin filaments (**Fig. 5C**). By adding increasing amounts of talin-VBS1 in the polymerisation assays, we showed that the 5M variant has a significantly decreased half maximal effective concentration, EC50 (**Fig. 5C and D**, and **movies S3 to S5**). Altogether these results suggest that while the K944A, R945A, D1013A, E1015A mutations yielded a slight increase in the affinity for actin filaments, the additional mutation at E986A position transforms vinculin into a more efficient actin bundler. Since vinculin activation by VBS1 is the rate-limiting reaction for vinculin-mediated bundling, our joint simulation and in vitro data suggest that the mutations in vinculin render Vt more exposed and accessible for actin by diminishing the stabilizing salt-bridges at the Vt-Vh interface just as VBS binding allosterically does (compare **Fig. 4B,C**).

**Figure 5:**
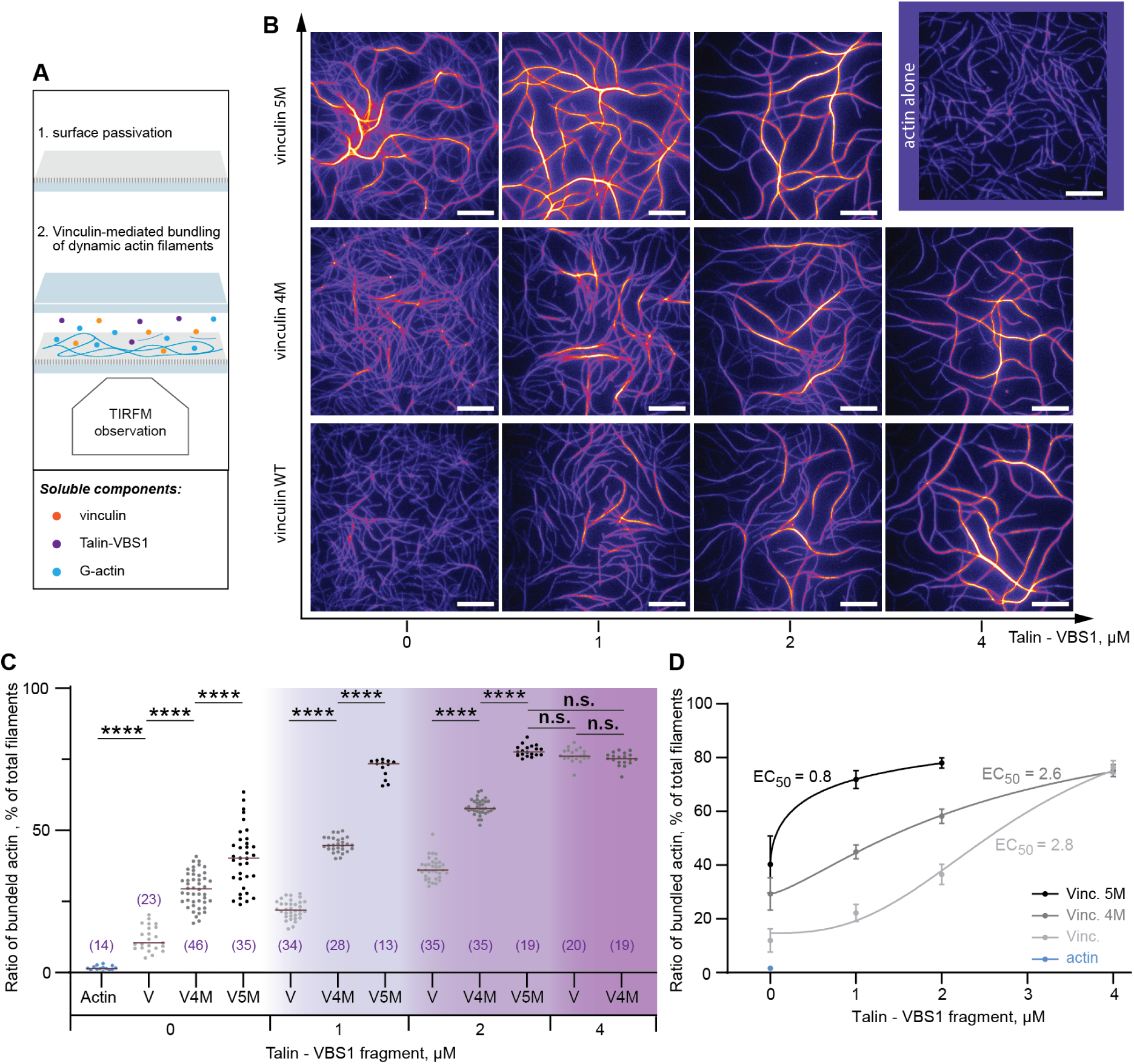
The 5M vinculin mutant forms stable actin filament bundles in the absence of talin. **A.** Schematic illustration of the experimental setup. (1) The passivated slides and coverslips were utilized to fabricate a chamber where (2) soluble proteins were injected at the onset of polymerisation / bundling reaction, as revealed by TIRFM. **B.** Representative images after 1 hour polymerisation of 0.6 µM Alexa-647 labelled actin alone, or in the presence 0.5 µM vinculin variants, or in the presence of 0.35 µM vinculin variants and 1, 2, or 4 µM talin VBS1. Scale bars, 10 µm. **C.** The relative amount of actin bundles was shown, as a ratio between bundled actin and the total filament population. Statistical comparisons using the Holm-Šídák test, and a one-way analysis of variance (ANOVA) showed significant variations among vinculin variants for their ability to generate actin bundles, with p values were <0.0001. **D.** Counts in **C** were plotted as a function of talin-VBS1 concentrations and fitted with a four-parameter dose-response equation (Graphpad software). The best fit of the overall dataset was obtained for 90% maximal bundle ratio and showed significant difference in the half maximal effective concentration, EC50. R^2^ ranged between 0.84 - 0.96, and was 0.94 for the overall dataset. Error bars represent standard deviation.

### Differential distribution of focal adhesions in vinculin-null MEFs following transfection with WT and mutant vinculin

Having established that the designed mutants mimic talin-activated vinculin in MT experiments and actin bundling assays, we next assessed the differential effects of WT and mutant (4M and 5M) vinculin forms on the formation and localization of cell-matrix adhesions in live cells. We transfected vinculin-null mouse embryonic fibroblasts (MEFs) with plasmids encoding the three molecules, tagged with GFP, to visualize the FA distribution in the transfected cells. Furthermore, the cells were fluorescently labelled for b1 integrin and F-actin (**Fig. 6A**). Naturally, the non-transfected (Control) cells were GFP-negative, yet they displayed poorly-organized b1 integrin, lacking the typical morphology of FAs. It is noteworthy that the control, un-transfected cells did form FA-like structures that contain other adhesome components, such as zyxin, as shown in **Fig. S7**. Moreover, F-actin distribution in these cells was rather diffuse throughout the cytoplasm.

**Figure 6:**
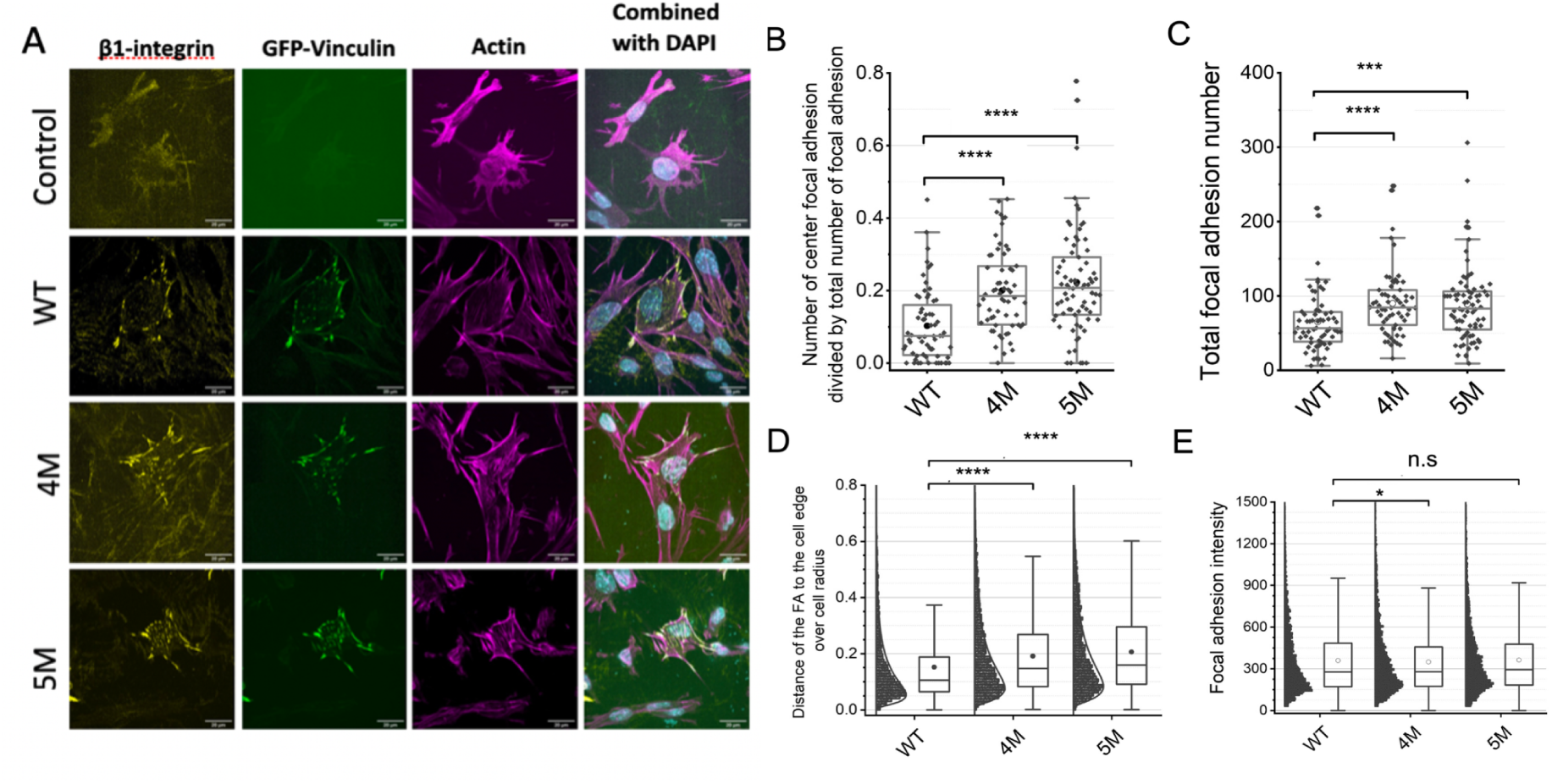
Vinculin mutants mimicking VBS binding form centrally-located adhesions distant from the cell edge. A) Images of MEF-vinculin-Null cells non transfected, or transfected with GFP-wild-type vinculin, GFP-4M, or GFP-5M vinculin mutants. Cells were stained with b-1 integrin antibody, phalloidin and DAPI. B) The fractions of center focal adhesion divided by the total number of focal adhesions of the cells. C) Quantification of total number of focal adhesions per cell. D) Quantification of the shortest distance of each focal adhesion to cell edge normalized to the equivalent cell radius. **E)** Quantification of focal adhesion intensity normalized to the background intensity. Box and whisker plotting: The box includes upper and lower quartile. The lower and upper whisker represents the lower quartile - 1.5 *interquartile range and upper quartile + 1.5 *interquartile range, respectively. The line in the box represents the medium and the dots represent the mean. Significant tests were performed using t-tests.

Expression of GFP-tagged WT vinculin induced a major reorganization of FAs and actin bundles, manifested by the formation of characteristic FAs, localized primarily at the cell periphery, and apparently recruiting b1 integrin into these sites. Expression of the mutant vinculin forms showed an enhanced capacity to form FAs, manifested mainly by the formation of multiple integrin adhesions that are located at the cell centre.

To quantify these differences in FA formation, we segmented the vinculin adhesions in 62-75 representative cells (**Table S2**) and defined those that are located at the cell centre and those located at the cells’ periphery (for details, see Materials and Methods). As shown in **Figure 6B and D**, the relative prominence of centrally-located adhesions in the cells expressing the mutant vinculins (both 4M and 5M) was significantly higher than in cells expressing WT vinculin. Furthermore, the average number of adhesions in cells expressing the mutant forms was ∼ 1.4 larger compared to WT (**Fig. 6C**). Interestingly, the fluorescence intensity of FA was essentially the same in the three experimental groups (**Fig. 6E**).

To further explore the possible mechanism underlying the differential properties of “central” and “peripheral” FAs, we transfected the vinculin-null MEFs with WT and mutant vinculin forms that are not tagged by GFP, and then immunolabeled the cells for vinculin and zyxin, in addition to phalloidin and DAPI. The choice of zyxin, whose localization in FAs was shown to be force-dependent^38, 47^ was motivated by the assumption that peripheral FAs are exposed to higher shear stress than central adhesions (see e.g. ref. ^48^). The images presented in **Fig. S7A** further support the results obtained with the GFP-tagged WT and mutant vinculins, namely that WT vinculin supports the development of peripheral FAs only, while the mutant forms support also the formation of prominent central adhesions. Notably, zyxin showed an exclusive association with the peripheral adhesions (even in cells expressing the 4M and 5M mutants), and was virtually absent from the central ones, which suggest that the low shear adhesions in the cell centre, that bind activated vinculin and not zyxin, might be fibrillar adhesions (see discussion section below). A quantification of the differential recruitment of the WT and mutant forms of vinculin is shown in **Fig. S7B**. Our observation supports the possibility that the mutations that were proposed to mimic the conformational changes that are physiologically driven by VBS1-mediated interaction of vinculin with talin indeed enable the mutated vinculin forms to bind and possibly support the lower-shear central FAs, that cannot recruit zyxin and WT vinculin.

## Discussion

Vinculin is a large multi-domain protein and a central hub of FAs and adherens-type cell-cell junctions that binds to the exposed vinculin-binding sites (VBS) of talin and alpha-actinin. Force activates both talin and vinculin, and both proteins activate each other but the molecular pathways of these force-dependent mutual allostery has remained poorly understood. We here lay out a comprehensive atomistic mechanism of the reciprocal talin-vinculin activation process, which we support by single-molecule experiments, actin bundling assays, and live cell based experiments, using wild-type and prospectively constitutively activated mutant vinculin.

Using MD simulation, we identified here a network of key salt-bridges at the head-tail interface of vinculin which stall the protein in the auto-inhibited closed conformation. A rewiring of this network upon VBS binding then loosens the closed state and facilitates an unfolding of the molecule. While previous MD simulations on short time scales of several hundred of picoseconds could not reveal an effect of bound VBS on the force response of vinculin^30^, our hundreds of nanosecond-long simulations are the first - to our knowledge - that resolve the activating effect of VBS at atomistic detail. Given the high structural and sequence conservation across different VBS, we propose this two-way allostery at the VBS-vinculin binding sites to be evolutionarily conserved and to also include non-talin VBS^19^.

A large body of work has confirmed the need of talin to be activated for vinculin by VBS exposure. A direct influence of force on vinculin activity has also been proposed but likewise questioned^31^. Our joint data underline that vinculin does not require to be subjected to force for binding talin. Instead, vinculin gets opened and activated upon talin binding in a largely force-independent manner, through the intricate maturation mechanism we discovered. As a consequence, we find the designed vinculin mutant mimicking talin binding, to localize to ‘low-shear’ central adhesions in cells. Previous studies (e.g. ^6^) demonstrated that cells spreading of fibronectin via ɑ5β1 integrin develop focal adhesion throughout their entire ventral membrane (mostly “central adhesions”), while cells adhering to vitronectin, via the ɑvβ3 receptors, form primarily “peripheral focal adhesions”. Here, expression of wild-type vinculin rescues β1 integrin engagement in peripheral focal adhesions (along with zyxin), while expression of mutant vinculin in these cells leads to association of β1 integrin in both peripheral and central adhesion sites. These novel observations support the notion that vinculin plays a key role in the recruitment of β1 integrins (most likely ɑ5β1) to adhesion sites (both peripheral and central) while zyxin association with focal adhesion is largely vinculin independent, and highly force depending.

Actomyosin-dependent forces, acting on vinculin are not required but can, of course, further enhance the maturation by shifting the vinculin conformational ensemble further to the extended open state of vinculin. In direct analogy, we predict the mechano-sensitive Focal Adhesion Kinase^49, 50^, which requires force to detach the auto-inhibitory from the kinase domain to be active only in the peripheral but not the central FAs. Instead, its constitutively active mutant Y180A/M183A should be able to phosphorylate its downstream substrates even in the low-shear central FAs, enhancing FA assembly also in the center of cells.

Our single-molecule force spectroscopy experiments have provided us with unprecedented quantitative insights into the force-dependent mutual regulation between talin and vinculin. Remarkably, the binding rates for VBS binding to full-length vinculin and D1 alone (or full-length vinculin head) are alike, indicating that only the D1 domain plays a role in recognizing and interacting with the stretched VBS and that talin binding is ignorant about the vinculin activation state. Our evidence seems to contradict recent work on the interaction between vinculin and the talin R6 domain, which showed a much decreased binding probability for FLV compared to that of Vd1^36^ (note that in contrast to this previous study using vinculin from bacterial expression, we here used eukaryotic expression with post-translational modifications in place). Our insights reveal the complex role of force in mutually regulating the interaction between vinculin and talin. While we previously showed how the force across talin directly controls vinculin binding—playing a biphasic role that establishes an optimal binding regime—, we demonstrate here that vinculin activation is regulated by a force-dependent allosteric mechanism. At forces below 10 pN, talin unfolds, exposing the VBS, and allowing vinculin binding. The interaction between the vinculin D1 domain and the mechanically stretched VBS weakens the head-tail vinculin interface, driving vinculin to a partially open or pre-active state, probably predisposed to actin recruitment. However, if talin unfolds at higher forces (>10 pN), vinculin quickly dissociates before evolving to the pre-active state (**Fig. 3F**), which could prevent actin recruitment. While the interaction between talin and active vinculin can resist high forces as those that can be transiently reached in a focal adhesion^10^, the binding event must occur at forces below 10 pN to allow for vinculin activation, and likely actin recruitment. Our model for the force-dependent allosteric regulation of vinculin activation establishes a much narrower force-range for optimal vinculin binding (5-10 pN versus the broader range 7-25 pN measured for D1), in better agreement with the average forces encountered in focal adhesions^10^.

Our joint scale-bridging data puts forward a novel force-dependent allosteric regulation that renders the talin-vinculin force-sensitive with a very distinct bimodal response. At low forces (up to ∼10 pN), talin-vinculin binding is sufficiently long to reach the “mature” state (activated or partially activated vinculin). Under higher forces (10-20 pN), vinculin-talin binding is short-lived as unbinding is faster than maturation, so the complex is very ephemeral. Thus, binding has to occur at a low range of forces for vinculin activation, a range largely overlapping with the physiological force range of 7-10 pN^9^. After maturation, forces on talin can increase up to ∼20 pN without compromising vinculin binding as the active mature state has been already reached.

The rationally designed vinculin mutants 4M and 5M lack the maturation and instead functionally and constitutively mimic vinculin with VBS being bound. The results from actin bundling and cellular adhesion assays confirm the mutants to act as talin-bound vinculin molecules, with the 5M mutant showing even stronger actin bundling capacity. The conservation of our mechanism across VBS observed in MD simulations suggests that the 4M and 5M vinculin mutants likely also serve as valuable mimics for other VBS-vinculin complexes such as those with alpha-actinin at cell-cell junctions. In fact, we observe VBS1 (and VBS3) to effect the additional mutation introduced in 5M more strongly than alpha-actinin, which could imply 5M to rather mimic the comparably strong binding of VBS1, while 4M rather captures vinculin binding to alpha-actinin. Overall, we propose 4M and 5M as versatile molecular tools to further dissect vinculin-mediated cellular mechanotransduction.

## Materials and methods

### Molecular Dynamics Simulations - Vinculin activation

The GROMACS^51^ 2018.4 simulation package was used for all simulations discussed in this study. The AMBER-ff99sb-ILDN force field^52^ with Joung ions^53^, and the Tip3p water model^54^ were utilized. Prior to production runs, the proteins were solvated, neutralized with a 0.15 M concentration of NaCl, and subjected to a 15 ps NVT and a 100 ps NPT equilibration. The NPT ensemble was kept at 300 K using the v-rescale thermostat^55^ and pressure was controlled at 1 atm using the Parrinello-Rahman barostat^56^ with a relaxation time of 2 ps. The use of hydrogen virtual sites^57^ allowed for an integration time step of 5 fs. The 1TR2 full-length vinculin structure^23^ excludes the proline-rich linker between residues P843 and P877. For our study, we inferred its conformation using the Chimera^58^ interface to MODELLER^59–61^.

To model the full-length VBS complexes, we used the pdb structures 1T01 (VBS1)^22^, 1RKC (VBS3)^23^, and 1YDI (ACT)^20^ and combined them with the full-length vinculin structure 1TR2 adapting the procedure described previously.^30^ Using pymol, we generated a hybrid structure minimizing the RMSD between the Cα-atoms of the D1 domains. The resulting structures were energy minimized in vacuum after which the simulation box was filled with Tip3p water and 0.15 M NaCl and the solvent energy was minimized by enforcing position restraints on the protein backbone and sidechains which were gradually released in a three-step procedure in an NVT ensemble. Finally, an extensive 1 microsecond-long NPT equilibrium simulation was conducted to ensure proper relaxation of the complexes.

For the force-probe MD runs, force was applied to the centre of mass of the tail domain, and on the the talin binding site located on the D1 domain.^23^ The apo-state recovery simulation were initiated from the resulting equilibrium conformation and, after removal of the VBS peptide, underwent the same energy minimization and equilibration scheme.

### Molecular Dynamics Simulations - Peptide extension

Here, the GROMACS^51^ version 2020.3 was used with the AMBER-ff99sb-ILDN force field with Joung ions, [62, 143] and the Tip3p water model. The simulation procedure was started with the VBS-D1 complexes structures with pdb-ID 1T01 (VBS1)^22^. Hydrogen virtual sites^57^ allowed an increase of the integration time step to 5 fs in the production set of simulations. Temperature and pressure were controlled as described above. Prior to production runs, the complex of VBS and vinculin head was equilibrated for 100 ns. For the production runs, we used distance-pulling along the x-axis applying a constant force to the N-terminal and C-terminal C-α atoms of the respective vinculin binding site. The VBS-vinculin-head complex we simulated 20×1 μs-long runs for 4 different forces. In case of the full length vinculin complex, 20×50 ns were simulated for 5 different forces.

### Single-Molecule Magnetic Tweezers Experiments

The single-molecule magnetic tweezers experiments were conducted on a custom-made setup, as previously described^42^. Experiments are carried out in custom-made fluid chambers composed by two cover slides sandwiched between a laser-cut parafilm pattern. The bottom glasses are cleaned by sonication on a three-step protocol: i) 1% Hellmanex at 50C, 30 min; ii) acetone (30 min); iii) ethanol (30 min), and then activated by plasma cleaning for 20 min to be silanized (immersion in 0.1% v/v 3-(aminopropyl)trimethoxysilane in ethanol for 30 min). Finally, the bottom cover slides are cured in the oven at 100 C for >30 min. The top cover slides are cleaned on 1% Hellmanex at 50 C, 30 min, and immersed in a repel silane (Cytiva, PlusOne Repel-Silane) for 30 min. The fluid chambers are assembled by melting the parafilm intercalator in a hot plate at 100C to ensure adhesion. After assembly, the chambers are incubated in a glutaldehyde solution (glutaraldehyde grade I 70%, 0.001% v/v) for 1 h, followed by incubation with 0.002% w/v polystyrene beads (∼2.5 µm diameter, Spherotech) for 20 min, and incubated with a 20 µg/ml solution of the HaloTag amine (O4) ligand (Promega) overnight. Finally, the chambers are passivated with a BSA-sulfhydryl blocked buffer (20 mM Tris-HCL pH 7.3, 150 mM NaCl, 2 mM MgCl2, 1% w/v sulfhydryl blocked BSA) for >3 h. The R3^IVVI^ polyprotein construct is freshly diluted in PBS at ∼2.5 nM and incubated for ∼30 min to achieve HaloTag binding. Finally, streptavidin-coated superparamagnetic beads (Dynabeads M270, Invitrogen) are added to bind with the biotinylated polyprotein terminus for (∼1 min). All experiments are performed on PBS buffer with 10 mM ascorbic acid (pH 7.3). After a talin molecule is found, full-length human vinculin is added to the fluid chamber by doing a buffer exchange to a buffer containing vinculin at the desired concentration at on 10 mM ascorbic acid.

### Expression and Purification of the Magnetic Tweezers Polyproteins

The (Ig32)_2_-(R3^IVVI^)-(Ig32)_2_ polyprotein was engineered by restriction digestion using the compatible cohesive and restriction enzymes BamHI and BglII between BamHI and KpnI sites. pFN18a (Ig32)_2_-(R3^IVVI^)-(Ig32)_2_ was subcloned into a modified pFN18a vector engineered with the AviTag^TM^ (Avidity) (sequence GLNDIFEAQKIEWHE). Constructs were cloned between the HaloTag at the N-terminal and the 6 histidine tag next to the AviTag^TM^ at the C-terminal. Recombinant plasmids were transformed in XL1Blue (Agilent Technologies) or Top10 (Thermofisher scientific) competent cells.

Polyprotein constructs were expressed in E. Coli BLR(D3) cells (Novagen). Cells were grown in LB supplemented with 100 mg/ml ampicillin at 37C. After reaching an OD_600_ of 0.6, cultures were induced with 1 mM of IPTG and grown at 25C for 16 hours. Cells were resuspended in 50 mM NaPi at pH 7.0 with 300 mM NaCl, 10% glycerol and 1 mM of DTT supplemented with 100 mg/ml lysozyme, 5 g/ml DNase, 5 g/ml RNase and 10 mM MgCl2 and incubated on ice for 30 minutes. This procedure was followed by gel filtration using a Superdex 200 10/300 GL column (GE Bioscienes). Proteins were stored in gel filtration buffer 10 mM HEPES pH 7.2, 150 mN. NaCl, 10% glycerol and 1 mM EDTA at -80C.

Gel filtration fractions were pooled and concentrated using Amicron® filters with selected MWCO. Constructs were biotinylated using BirA Biotin Ligase (Avidity) following the manufactures suggested protocol. Biotinylation was confirmed using Streptavidin HRP conjugate (Millipore) with biotinylated/unbiotinulated MBP-AviTag^TM^ fusion protein (Avidity) as controls.

### Human Vinculin mutagenesis for bacterial protein expression

Introduction of mutations into human vinculin (termed here M4 & M5) was based on the modeling described in **Fig. 4A, B** above. The mutations were introduced into a bacterial expression vector pBXNHG3_huVinculin (provided by Rajaa Boujemaa and Ohad Medalia, Univ. of Zurich) containing an N-terminal 10His tag and sfGFP-3C cleavage site. The mutations were introduced into the huVinculin coding sequence by the TPCR method^62^ using Q5 High-Fidelity Master Mix (NEB). In the initial step the 4 mutations (K944A_R945A_D1013A_E1015A) were introduced, and the 5^th^ (E986A) mutation was then added. The primers that were used to introduce the mutations are:

Vinculin_944A,945A_Forward:

GAGGGGGCAGTGGTACCgctgcaGCACTCATTCAGTGTGCC

Vinculin_1013A,1015A_Reversed (used with the two forward primers):

CTCTGTGGCCTGCTCAGAtgCCTCAgCACTGATGTTGGTCCGGC

Vinculin_986A_Forward:

CAACCTCTTACAGGTATGTGccCGAATCCCAACCATAAGCACC

The entire ORF, including the promoter region was subsequently verified by DNA sequencing.

### Human full-length vinculin expression in insect cells

Full-length vinculin variants (wild type, V4M and V5M) cDNAs were amplified (using the forward 5’-ATATATGCTCTTCTAGTATGCCAGTGTTTC-3’, and the reverse 5’-TATATAGCTCTTCATGCCTGGTACCAGGG-3’ primers) of 1066 amino acids long, was cloned into a pFBXNH3 insect cell expression vector containing an N-terminal His tag followed by a 3C protease cleavage site, using the fragment exchange (FX) cloning strategy (Geertsma and Dutzler, 2011). Protein expression was performed by generating a recombinant baculovirus for ExpiSf9 insect cell infection at a density of 2.0×10^6^ ml^-1^ using the Bac-to-Bac system (Invitrogen). Briefly, the recombinant vinculin encoding plasmid pFBXNH3 was transformed into DH10Bac E. coli cells, which enabled the transposition of the recombinant gene into the bacmid genome. The recombinant bacmid DNA was then isolated and transfected into ExpiSf9 cells to generate the full-length vinculin-expressing baculovirus.

### Protein expression, purification and fluorescence labelling

ExpiSf9 cells were infected with the full-length vinculin-expressing baculovirus for recombinant protein expression as described^28^. Briefly, expressing cells were harvested by gentle centrifugation and lysed by sonication in 20 mM Tris-HCl pH 7.5, 0.4 M NaCl, 5% Glycerol, 1 mM DTT, 0.1% Triton, and protease inhibitors. Full-length vinculin variants were purified from clarified cell extract using a Ni-Sepharose 6 Fast Flow metal affinity chromatography, HisTrapFF (Cytiva Lifesciences). Proteins were further purified on a Superdex 200 size exclusion column (Cytiva Lifesciences) and eluted with 20 mM Tris-HCl pH 7.8, 0.15 M KCl, 1 mM MgCl2 and 5% Glycerol. Protein aliquots were supplemented with 50% glycerol and stored at -20°C.

Actin was purified from rabbit skeletal muscle acetone powder, purified according to the method of Spudich and Watt^63^, and stored in G-buffer (5 mM Tris-HCl pH 7.8, 0.2 mM CaCl2, 0.5 mM DTT, 0.2 mM ATP). Actin was further labeled on cysteine with Alexa-Fluor 647 maleimide dyes (Invitrogen). Labeling was performed in 2 mM Tris HCl pH 7, 0.2 mM CaCl_2_, 0.2 mM ATP, for 16h. To limit excessive labeling, protein/dye molar ratio was kept ≤ 1:3. The reactions were stopped by adding 10 mM DTT. Labeled actin was polymerized, depolymerized, and gel-filtered using Superdex 200 size exclusion column (Cytiva Lifesciences), in 2 mM Tris HCl pH 7, 0.2 mM CaCl_2_, 0.2 mM ATP, 0.5 mM DTT. Ca-monomeric actin was stored on ice and used within 2 weeks for TIRFM. Human Talin 1 VBS1 residues 482-636 construct was expressed in BL21 (DE3)pLysS strain and purified using using a Ni-Sepharose 6 Fast Flow metal affinity chromatography as described in ref. ^28^, supplemented with 20% glycerol, aliquoted, flash-frozen in liquid nitrogen and stored at -80°C.

### Glass surface passivation

Slides and coverslips (CVs) were used to assemble the reaction chambers were drastically cleaned by successive chemical treatments, and coated with tri-ethoxy-silane-PEG (5 kDa, PLS-2011, Creative PEGWorks, USA) 1mg/ml in ethanol 96% and 0.02% of HCl, as described previously.^28^ mPEG-silane coated slides and CVs were then stored in a clean container and used within a week time.

### TIRFM imaging

Reconstitution assays were performed using fresh actin polymerization buffer, containing 20 mM Hepes pH 7.0, 40 mM KCl, 1 mM MgCl2, 1 mM EGTA, 100 mM β-mercaptoethanol, 1.2 mM ATP, 20 mM glucose, 40 μg/mL catalase, 100 μg/mL glucose oxidase, and 0.4% methylcellulose. The final actin concentrations were 0.6 µM, with 20% Alexa monomer labels. The polymerization medium was supplemented with vinculin variants and talin VBS1, as indicated in the figure legends and methods. The reaction medium was rapidly injected into a passivated flow cell at the onset of actin assembly, imaging started after 2 min. Time-lapse TIRFM was recorded every 15 sec. TIRF images were acquired using a Widefield/TIRF – Leica SR GSD 3D microscope, consisting of an inverted widefield microscope (Leica DMI6000B / AM TIRF MC) equipped with a 160x objective (HCX PL APO for GSD/TIRF, NA 1.43), a Leica SuMo Stage, a PIFOC piezo nanofocusing system (Physik Instrumente, Germany) to minimize the drift for an accurate imaging, and combined with an Andor iXon Ultra 897 EMCCD camera (Andor, Oxford Instruments). Fluorescent proteins were excited using a solid-state diode laser 642 nm (500 mW). Laser power was set to 3% and dyes were excited for 50 ms. Image acquisition was performed with 25 degrees-equilibrated samples and microscope stage. The microscope and devices were driven by Leica LAS X software (Leica Microsystems, GmbH, Germany).

### Image processing and data analysis of fluorescence images

Time-lapse videos of filament growth and steady state images taken after 1 hour of actin assembly were processed with Fiji software (NIH). 15 to 35 steady state images were taken for each condition. To determine the ratio of bundled actin to the total assembled filaments, macros written in Fiji allowed to first subtract the background for each individual image, then average the min and max fluorescence intensities for single actin filaments, bin grey values (256 per 65535 grey levels) and calculate histogram counts for individual images. This allowed to evaluate counts of pixels occupied by single actin filaments or by bundled actin.

### Human Vinculin mutagenesis for mammalian expression

The bacterial Vinculin mutant constructs were served as templates for PCR amplification of 10His-GFP-3C-Vinculin K944A_R945A_D1013A_E1015A (huVin_M4) or K944A_R945A_D1013A_E1015A E986A (huVin_M5) mutant sequences. The resulting PCR fragments were cloned into pCDNA3.4 by restriction enzymes and ligation. Using different sets of primers but the same bacterial vinculin mutant constructs as templates. 10His-Vinculin mutant sequences, without GFP-3C sequences were PCR amplified and cloned into pCDNA3.4 by restriction enzymes and ligation. The WT Vinculin sequence was cloned similarly by restriction enzyme and ligation to obtain 10His-Vinculinwt_pCDNA3.4 construct.

### Cell culture

Vinculin-null mouse embryonic fibroblasts (Vin-null-MEFs), derived from vinculin-null mice, were generously provided by Eileen Adamson (Burnham Institute)^64^. The cells were grown in DMEM (Gibco, Grand Island, New York) containing 10% FCS and 100 U/mL PenStrep (Biological Industries, Beit Haemek, Israel) at 37° C in a 5% CO2 humidified atmosphere.

### Cell transfection and immunofluorescence microscopy

#### Expression of GFP-tagged vinculin variants (WT and 4M/5M mutants) in Vin-null MEFs

The vinculin-null MEFs were seeded onto optical glass-bottom 96-well plates (Cat#164588 Thermo Scientific Nunc) coated with 10µg/ml Bovine fibronectin (Biological Industries –Currently Sartorius). After 24 hours, the cells were transfected with the different vinculin variants, tagged with GFP, using TurboFect (Thermo Fisher Scientific, Waltham, MA) as a transfection agent. The transfection was conducted according to the manufacturers’ protocols. After 24h, the cells were simultaneously permeabilized and fixed, with 0.5% Triton X100 in 3% paraformaldehyde for 2 minutes and further fixed with 3% paraformaldehyde for additional 30 minutes. Then, the cells were stained with beta-1 integrin antibody (HUTS21, BD Bio-sciences) or mouse anti-human β1 integrin P5D2 (Developmental studies Hybridoma bank)”, Phalloidin TRITC (Sigma) and DAPI (Sigma).

#### Expression of His-tagged vinculin variants in vinculin-null MEFs

His-tagged vinculin variants (WT and the 4M/5M mutants) were transfected into Vin-null- MEFs, and the cells were processed as above and co-stained for vinculin (monoclonal) and zyxin (polyclonal, both prepared by the Antibody Production Laboratory of the Department of Biological Services, Weizmann Institute of Science), Phalloidin TRITC (Sigma) and DAPI (Sigma).

### Imaging

Fluorescence images were acquired using a DeltaVision Elite system (GE Healthcare/ Applied Precision, USA), mounted on an inverted IX71 microscope (Olympus, Japan) and equipped with a Photometrics CoolSNAP HQ2 camera (Photometrics, Tucson, AZ). The system was running SoftWorX 6.1.3. Pictures were acquired using an Olympus UIS2 BFP1 60×1.42 PlanApoN oil objective (Olympus, Japan).

### Image analysis

Confocal images or wide-field fluorescent images of vinculin or zyxin were first segmented using Ilastik software (https://www.ilastik.org). After the segmentation, individual objects were identified, and the object with a size larger than 0.3 μm^2^ were preserved and considered as a “focal adhesion-like structure”. Confocal or wide-field fluorescent images of actin were used to identify cell boundaries. Intensity thresholding by Otsu’s method^65^ was applied to segment the cells. After the segmentation, the object with the largest area was preserved and considered as the main cell body. The approximate cell radius was then defined as

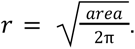

Quantification of FA area, length and fluorescence intensity was performed using a custom code written in Matlab (MathWorks, Natick, MA) after correcting for fluorescence bleaching.

### Focal adhesion intensity measurement

Since the intensity of focal adhesions and related structures can be influenced by the background intensity of the cell, especially in wide-field fluorescent images, we normalized the focal adhesion intensity by subtracting the background intensity. The local background intensity was calculated for each focal adhesion by measuring the average intensity of the area surrounding the focal adhesion. The area outside of the cell and other focal adhesions are excluded from the background intensity calculation. Average intensity of each focal adhesion area was then calculated after the subtraction of background intensity.

### Focal adhesion location measurement

The distance between a focal adhesion and the cell’s edge was obtained by calculating the shortest distance between the center of mass of the focal adhesion and the cell boundary. The “central focal adhesions (possibly – “fibrillar adhesions”) were defined as those adhesions whose distance from the cell edge is larger than 0.3×*r* (*approximate cell radius*).

## Acknowledgements

Acknowledgement

FG and BG thank the Klaus Tschira Foundation for generous financial support. The study was also supported by the Minerva Center on “Aging, from Physical Materials to Human Tissues” to B.G.; the Schweizerischer Nationalfonds zur Förderung der Wissenschaftlichen Forschung to O.M. (SNSF 310030_207453); the HGS MathComp graduate school of Heidelberg University, the Max Planck School Matter to Life supported by the German Federal Ministry of Education and Research (BMBF) in collaboration with the Max Planck Society, the Flagship Initiative funded by the Federal Ministry of Education and Research (BMBF) and the Ministry of Science Baden-Württemberg within the framework of the Excellence Strategy of the Federal and State Governments of Germany, the state of Baden-Württemberg through bwHPC and the German Research Foundation (DFG) through grant INST 35/1134-1 FUGG to F.G.. This work was supported in part by the Francis Crick Institute which receives its core funding from Cancer Research U.K. (FC001002), the U.K. Medical Research Council (FC001002), and the Wellcome Trust (FC001002). R.T-R. is recipient of a King’s Prize Fellowship. This work was supported by the European Commission (Mechanocontrol, Grant Agreement 731957), BBSRC sLoLa (BB/V003518/1), Leverhulme Trust Research Leadership Award RL 2016-015, Wellcome Trust Investigator Award 212218/Z/18/Z and Royal Society Wolfson Fellowship RSWF/R3/183006 to S.G.M.

