## Supplementary Information for "How talin allosterically activates vinculin"

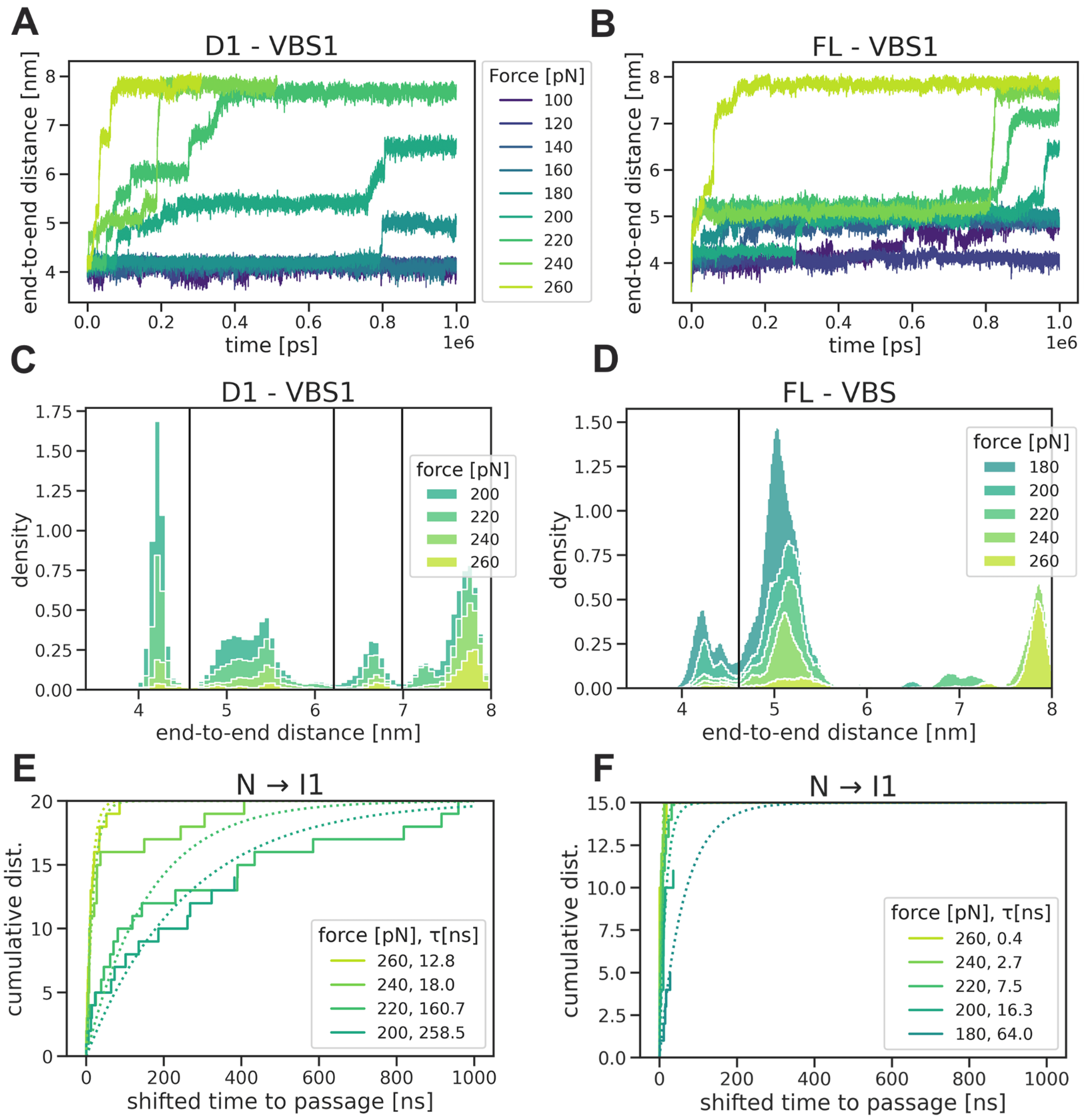


**Figure S1: Adopt from 4.4 top and 4.5 top Kinetic analysis of the force-probe MD simulations of VBS1 bound to D1. (A,B)** End-to-end distance traces for the pulled VBS1 peptides in complex with the Vinculin D1 domain (A) and the full protein (B) for forces between 100 and 260pN. **(C,D)** Stacked distributions of end-to-end lengths observed during 1microsecond long MD simulations at four different constant forces. The locations of the main barriers along the reaction coordinate, identified by a low density of end-to-end distances at that distance, are indicated by the black horizontal lines. The initial unfolding step from U (unfolded) to I1, the first intermediate, initiated the full unfolding that in experiments is known to result in dissociation. Thus the U->I1 transition was used as a proxy of vinculin-VBS dissociation. **(E,F)** For each force, the dwell time of the unfolded state N to I1 was recorded and shown is the cumulative distribution of these dwell times. Data was fit with a single exponential function to deduce the VBS unfolding time (denoted mean first passage time, MFPT, in the main text) for each given force (inset).

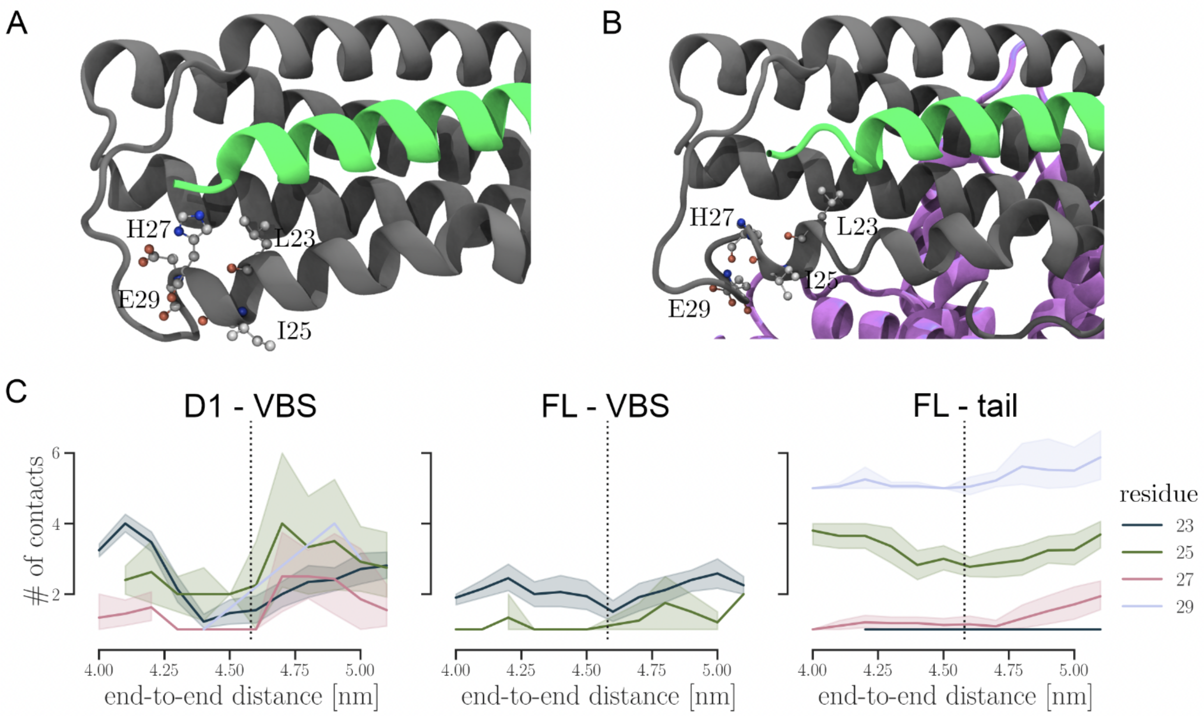


**Figure S2: Vinculin tail and talin VBS compete for the interaction with D1.** A) Cartoon representation of the vinculin D1 domain (grey) - VBS1 (green) interaction. Residue L23 and H27 reinforce the complex. B) With full-length vinculin, L25 and E27 orient towards the vinculin tail (purple). C) The evolution of selected inter-residue contacts with respect to the VBS end-to-end distance. Left) D1 residues in contact with VBS for the D1-VBS1 complex. Center) Residues in contact with VBS when simulating the full length vinculin complex. Right) The number of contacts between D1 and the tail domain. The solid lines show the average obtained from up to 60 simulations, the shaded area represents the standard deviation.

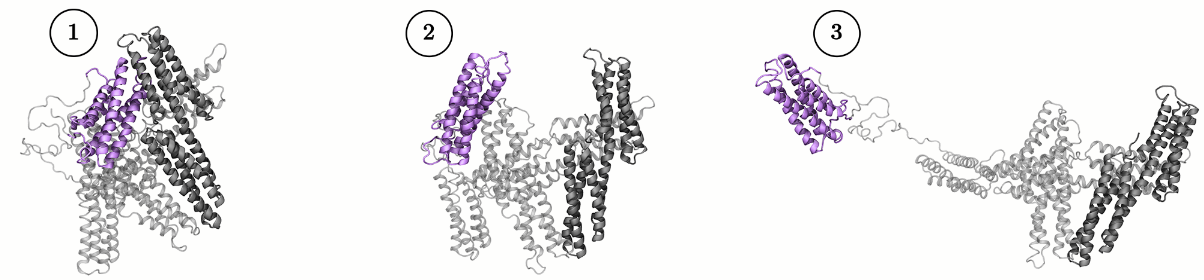


**Figure S3: Snapshots of head-tail dissociation in apo-vinculin as sampled by FPMD simulations.** Coloring as in Fig. 2A of main text.

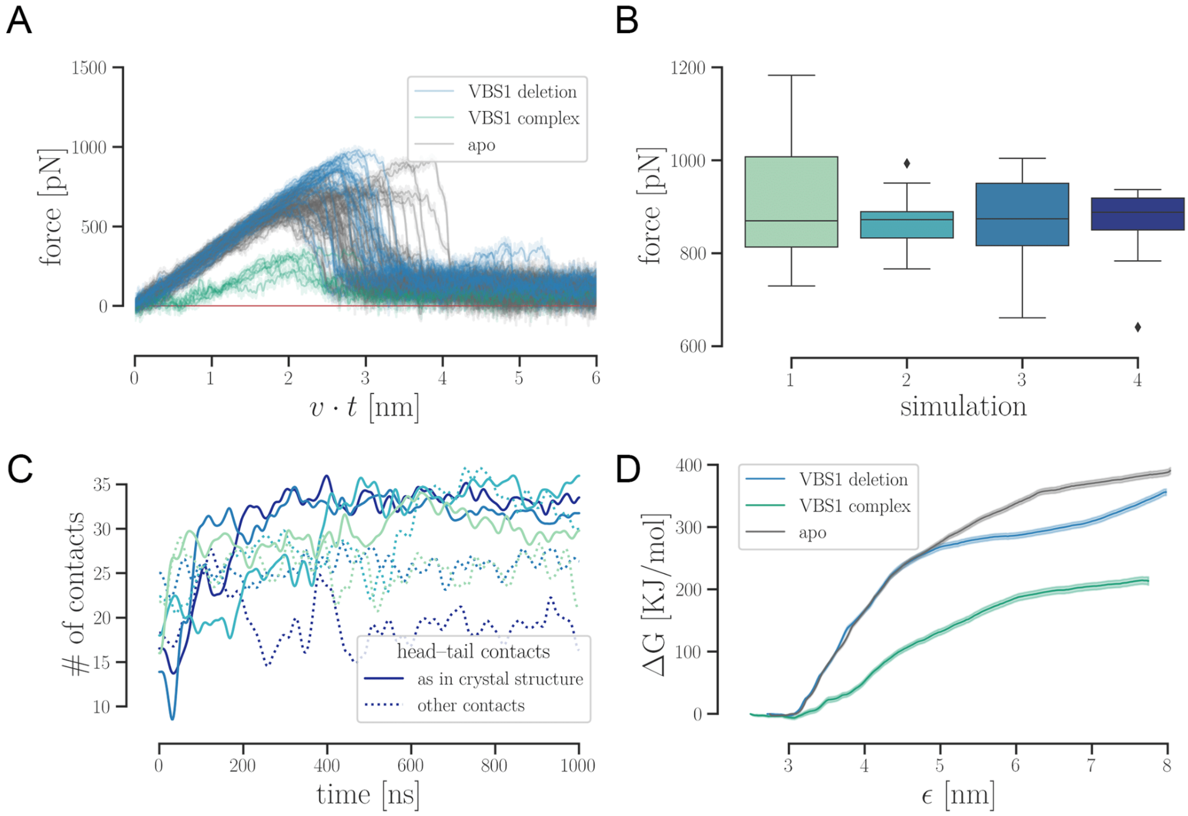


**Figure S4: Loosening of head-tail interface is caused by VBS binding and not the use of different starting crystal structures,** as shown by additional equilibrium, FPMD, simulations not started from the apo crystal structure but instead from the VBS-vinculin crystal structure, from which VBS was removed and vinculin equilibrated. **A:** Force extensions curves for force-probe simulations of the VBS1 complex (green), the apo-state protein (gray) and the 'recovered' apo state after the VBS1 peptide was deleted and the resulting structure was equilibrated. **B:** Following VBS1 deletion, four independent equilibration runs were carried out. The box plot shows the highest forces observed in 10 pulling simulations starting from each of the relaxed structures. **C:** The solid lines show the number of contacts that are recovered during the four equilibration simulations. As a reference, we used 53 head-tail contacts identified in the crystal structure with a 3.5 cut-off. Dotted lines show the number of non-apo contacts. **D:** Mean (solid line) and standard deviation (shaded area) of the free-energy profiles calculated by umbrella sampling for the opening of the described structures.

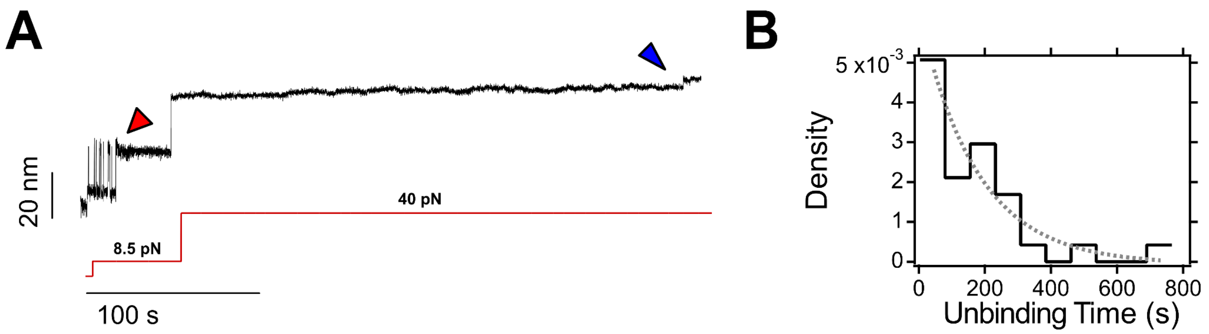


**Figure S5: Unbinding kinetics of full-length vinculin head. (A)** Typical magnetic tweezers recording of full-length vinculin head binding (read arrow) and unbinding (blue arrow) from talin $R3^{IVVI}$. At 20 nM vinculin head, binding at 8.5 pN occurs after a few seconds; however, to dissociate the complex at 40 pN requires a few hundreds of seconds, similar to the Vd1 domain, suggesting that of vinculin head, only the D1 domain participates in the talin-vinculin interaction. **(B)** Distribution of unbinding times of full-length vinculin head from $R3^{IVVI}$ at 40 pN. The unbinding kinetics are distributed as a single exponential, indicating a single bound-mode, similar to Vd1 and in contrast with full-length vinculin. Data from N=30 unbinding events.

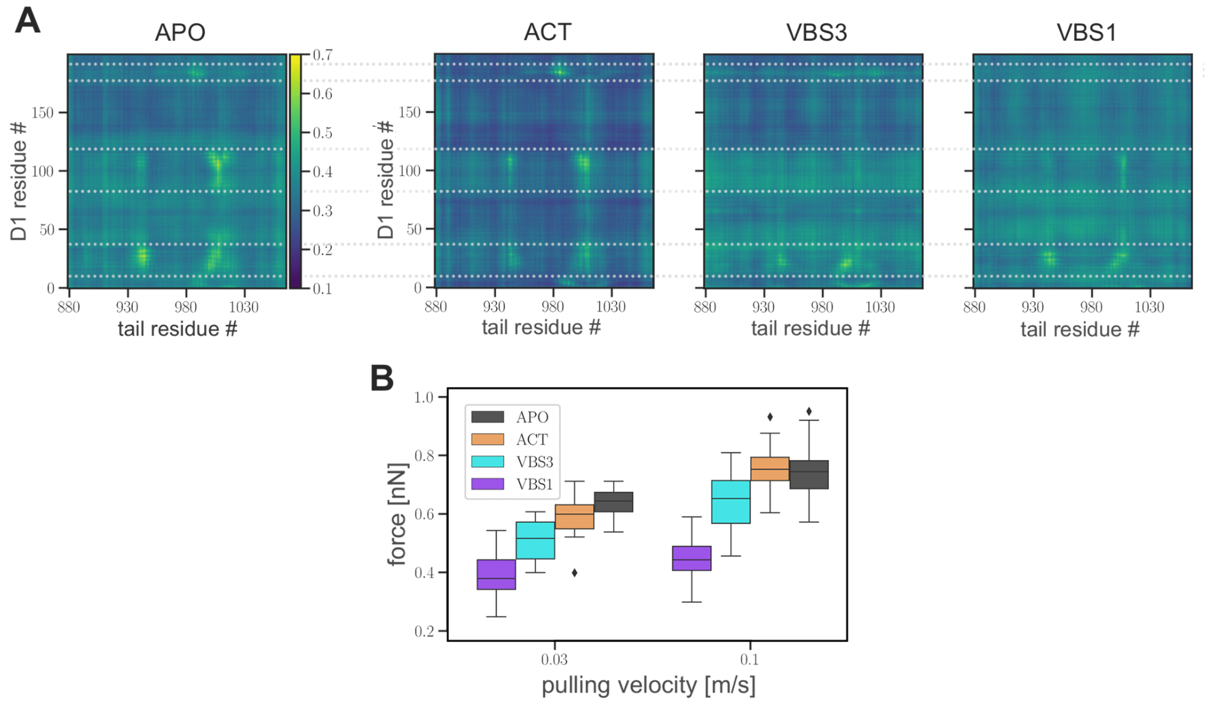


**Figure S6: Different VBS allosterically affect vinculin head-talin interactions differently. A**: The interactions between vinculin D1 and tail portrayed by generalized correlations maps. The data shown is extracted from force-probe MD trajectories with a pulling speed of 0.1m/s. The correlations were computed for each simulation using a 5 ns-long fraction of each trajectory that precedes the moment of 300 pN force across the protein. The average of 20 runs is presented for each complex type. **B**: Different VBSs dissimilarly impact the rupture force. The distributions of the highestm observed force in each trajectory for a set of simulations ranging over two pulling speeds (0.03m=s and 0.1m=s) are represented by a box plot. For each velocity and complex type, the plot represents data from at least 10 replicas.

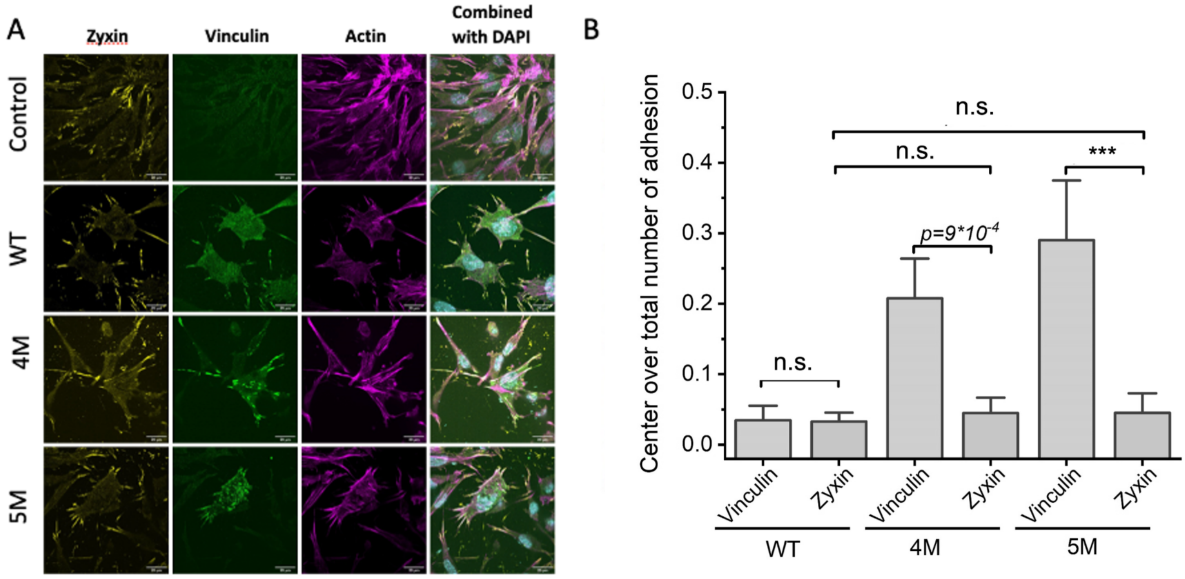


**Figure S7: 4M and 5M vinculin but not zyxin show a strong propensity to localize in central ("low shear") adhesions. A)** Images of MEF-vinculin-Null cells non-transfected or transfected with His-tag-wild-type vinculin, His-tag-4M, or His-tag-5M vinculin mutants. Cells were immune-stained with anti-Zyxin, anti-Vinculin antibodies, phalloidin and DAPI. **B)** Quantifications of center vinculin-positive focal adhesions over the total vinculin-positive number adhesions and center zyxin-positive focal adhesion over the total zyxin-positive number adhesions in MEF-vinculin-Null cells transfected with wild-type Vinculin, 4M or 5M Vinculin mutant.

| cluster | ID1 | ID2 | correlation loss |
|---:|---:|---:|---:|
| 0 | 185 | 987 | -0.279 |
| 1 | 113 | 1013 | -0.386 |
| 2 | 33 | 945 | -0.289 |
| 3 | 20 | 1013 | -0.262 |
| 4 | 113 | 1004 | -0.296 |
| 5 | 93 | 1011 | -0.247 |

**Table S1: Residue pairs with highest correlation loss.** The clusters identified in Fig. 4A were scanned for the strongest contributors which are summarized in this table and colored according to Fig. 4B.

| | Number of cells | Number of focal adhesions |
|---|---|---|
| WT | 64 | 4168 |
| 4M | 62 | 5580 |
| 5M | 75 | 6677 |

**Table S2: Number of cells and focal adhesions** used the quantification

**Movie S1 : Assembly of actin filaments.** Actin filament polymerization in the presence of 0.6 µM actin Alexa-647 labeled, followed by TIRFM. Note that the filaments drift and collide without fusing into bundles. Related Figure 5.

**Movie S2 : Vinculin 5M mediates stable bundling of actin filaments in the absence of VBS1.** Actin polymerization and bundling in the presence of 0.6 µM actin Alexa-647 labeled, 500 nM of the indicated vinculin variant, followed by TIRFM. Related Figure 5

**Movie S3 : Vinculin 5M induces increased bundling of actin filaments when supplemented with 1 µM VBS1.** Actin polymerization and bundling in the presence of 0.6 µM actin Alexa-647 labeled, 350 nM of the indicated vinculin variant, and 1 µM talin-VBS1, followed by TIRFM. Related Figure 5.

**Movie S4 : Vinculin 5M mediates higher degree of actin bundling in the presence of 2 µM VBS1.** Actin polymerization and bundling in the presence of 0.6 µM actin Alexa-647 labeled, 350 nM of each vinculin variant, and 2 µM talin-VBS1, followed by TIRFM. Related Figure 5.

**Movie S5 : Vinculin 5M activation by VBS1 is more efficient than vinculin 4M and the WT.** Actin polymerization and bundling in the presence of 0.6 µM actin Alexa-647 labeled, 350 nM of vinculin WT or V4M, and 4 µM talin-VBS1, while 350 nM of vinculin V5M with 2 µM talin-VBS1 yields thicker bundles, followed by TIRFM. Related Figure 5.
